# Noradrenaline prolongs the state of motor arrest via cerebrospinal fluid signaling

**DOI:** 10.64898/2026.02.06.704024

**Authors:** Mahalakshmi Dhanasekar, Kevin Fidelin, Mattia Greco, Elias T Lunsford, Martin Carbo-Tano, Charlotte Deleuze, Lionel Moisan, Thierry Mora, Aleksandra Walczak, Claire Wyart

**Affiliations:** Institut du Cerveau (ICM), Sorbonne Université, UPMC Univ Paris 06, Inserm, CNRS, AP-HP, Hôpital Pitié-Salpêtrière, 75013 Paris, France; Laboratoire de physique de l’École Normale supérieure, CNRS, PSL University, Sorbonne Université and Université de Paris, 75005 Paris, France; Department of Neuroscience, Faculty of Health and Medical Sciences, University of Copenhagen Blegdamsvej 3, 2200 Copenhagen, Denmark; Université Paris Cité, CNRS, MAP5, 75270 Paris Cedex 06, France

## Abstract

Neuromodulators can mediate whole-body physiological changes, however how signaling from relatively small populations of neurons produces widespread and sustained effects remains incompletely understood. Here, we demonstrate that sustained noradrenergic activation induces prolonged motor arrest accompanied by postural collapse, bradycardia, and emetic responses. Motor suppression arises from inhibition of brainstem commands to the spinal cord rather than motoneuron silencing. Noradrenaline induces a glial calcium wave that propagates throughout the central nervous system. This wave most prominently persists at the brainstem-spinal cord boundary and ventricular midline. At this location, ependymal radial glia express Adra1a receptors on cilia contacting cerebrospinal fluid (CSF). Ventricular noradrenaline injections recapitulated the glial wave and motor arrest, while CSF administration of an Adra1 antagonist shortened motor arrest. Altogether, noradrenaline sustains motor arrest by signaling through the CSF via ciliary Adra1a receptors on ependymal radial glia showing that cerebrospinal fluid as a critical transmission route for persistence of behavioral states.

**Highlights:**

- Noradrenergic surges coordinate a whole-body response with motor arrest, postural disruptions, bradycardia, and emesis-like reflexes
- Motor arrest results from silencing of brainstem motor commands to the spinal cord
- Noradrenaline-induced glial calcium waves last longest at the obex/ventricular midline where cilia of CSF-contacting glial cells bear Adra1 receptors
- Noradrenergic signaling in the CSF is necessary and sufficient for motor arrest to persist

## Introduction

Unlike fast synaptic transmission, which operates at millisecond timescales within defined synapses, neuromodulators act over seconds to minutes across distributed brainstem, spinal, and peripheral networks to establish coherent physiological states ^1–4^. One such state includes threat responses, which span a striking physiological spectrum, encompassing autonomic reactions like bradycardia^5–7^, nausea and the emetic reflex ^8^, and dramatic shifts in muscle tone—from paralytic freezing to full postural collapse ^9–11^. Such whole-body integration is critical for selecting the safest behavioral response to ensure survival and relies on central and peripheral mechanisms involving neuromodulators ^12^. Noradrenaline is particularly well-suited for this role: released centrally, it modulates arousal, vigilance, attention, and locomotion through synaptic and volume transmission^13–16^, peripherally it controls autonomic functions, including heart rate and blood pressure ^17,18^.Upon threat, aversive stimuli recruit noradrenergic neurons in the locus coerelus^19,20^ and in the medulla ^21^. Strong aversive stimuli trigger motor arrest, which lasts tens of seconds ^9,22–27^, and induce bradycardia^5,28,29^ Yet, how noradrenergic signaling modulates motor circuits to elicit motor arrest over such long timescales remains poorly understood.

Vertebrate locomotion relies on a hierarchical circuit architecture distributed throughout the central nervous system: higher brain centers, including basal ganglia and the mesencephalic locomotor region (MLR)^30–35^ project onto reticulospinal neurons^36^, which in turn recruit spinal central pattern generators (CPGs) ^36–39^. CPGs drive motor neurons to execute patterned muscle contractions^40–45^, with sensory feedback refining motor output throughout ^46–48^. Motor arrest involves sustained disruption of either motor circuit elements (higher motor centers, reticulospinal neurons, CPGs, or motor neurons) or communication between them. Recent studies have revealed that motor arrest is associated with a noradrenergic dependent glial calcium wave^49–51^ that elicits downstream adenosine signaling^51–53^. Yet which motor circuit elements are disrupted during motor arrest and how subsequent motor suppression lasts for long durations (10s of seconds) remain elusive.

To encompass both major questions mentioned above, we optogenetically activated noradrenergic neurons in larval zebrafish that induced sustained motor arrest together with autonomic defects and postural collapse. Reduction in synaptic drive to motor neurons resulted in motor arrest, which was followed by a refractory period lasting about 15 s, during which supraspinal commands could not induce motor output. This refractory period coincided with coincided with sustained noradrenaline-evoked glial calcium wave at the obex and midline, where ependymal cells expressing α1-adrenergic receptors on their cilia are exposed to the cerebrospinal fluid. Ventricular noradrenaline delivery recapitulated both the calcium wave and motor suppression, while an α1-adrenergic antagonist reduced it. These findings indicate that noradrenaline sustains motor arrest through cerebrospinal fluid signaling via Adra1a-expressing ependymal glia.

## Results

### Graded noradrenergic neuron activation induces a prolonged motor arrest with postural disruptions, bradycardia and emetic like reflexes

Noradrenergic neurons are typically recruited during aversive stimuli^19–21,50^. In zebrafish, *dbh+* noradrenergic neurons are primarily present around the obex. To determine whether sustained noradrenergic activation is sufficient to coordinate behavioral and autonomic responses, we optogenetically activated *dbh+* neurons in freely-swimming 6 days post fertilization (dpf) *Tg(dbh:KalTA4;UAS:CoChR-eGFP)* larval zebrafish. We exposed larvae to optogenetic trains of variable duration (0.25-3s, **Figure 1A**), consisting of 50 ms-long light pulses at 10Hz, mimicking phasic noradrenergic firing during salient stimuli ^13,54^. Loose-patch recordings confirmed that single 50 ms-long light pulses reliably evoked 1-3 spikes in noradrenergic neurons expressing CoChR-eGFP (**Figure S1A,B**). To eliminate visually induced motor responses to blue light stimulation, we performed enucleation at 3 dpf (**Figure S1C,D**). Trains lasting at least 0.5 s increased locomotion in control siblings, while they immediately suppressed locomotion in transgenic larvae (n = 16 per genotype, 4 trials per larva; **Figure 1B**, **Figure S1E**). From baseline, with inter bout intervals lasting 2-3 seconds (2.19 ± 0.32 s; 2.98 ± 0.61 s; 2.15 ± 0.29 s), motor arrest duration lasted 10.53 ± 3.57 s upon 0.5s long train stimulation, 37.25 ± 8.83 s upon 1s-long train stimulation, and 72.88 ± 14.82 s upon 3s-long train stimulation (**Figure 1C, S1D**), while 0.25 s stimulations showed no significant effect. In addition, stimulation trains longer than 1 s also induced postural disruptions characterized by rolling to the side (9.6 ± 4.5% at 1 s, 44.2 ± 7.0% at 3 s; **Figure 1D; supplementary video 1**) while control siblings failed to do so. Along with this postural loss, 3 s activation induced visceral motor responses reminiscent of emetic reflexes (**Figure 1E; supplementary video 2)** and bradycardia (**Figure 1F–H**; decrease of 0.80 ± 0.13 Hz occurring 4.25 ± 0.37 s after stimulation; n = 4 larvae, 8 trials). These results demonstrate that sustained noradrenergic activation is sufficient to trigger a coordinated whole-body response encompassing an immediate and long-lasting motor arrest, with postural disruption, visceral motor responses, and bradycardia.

**Figure 1.**
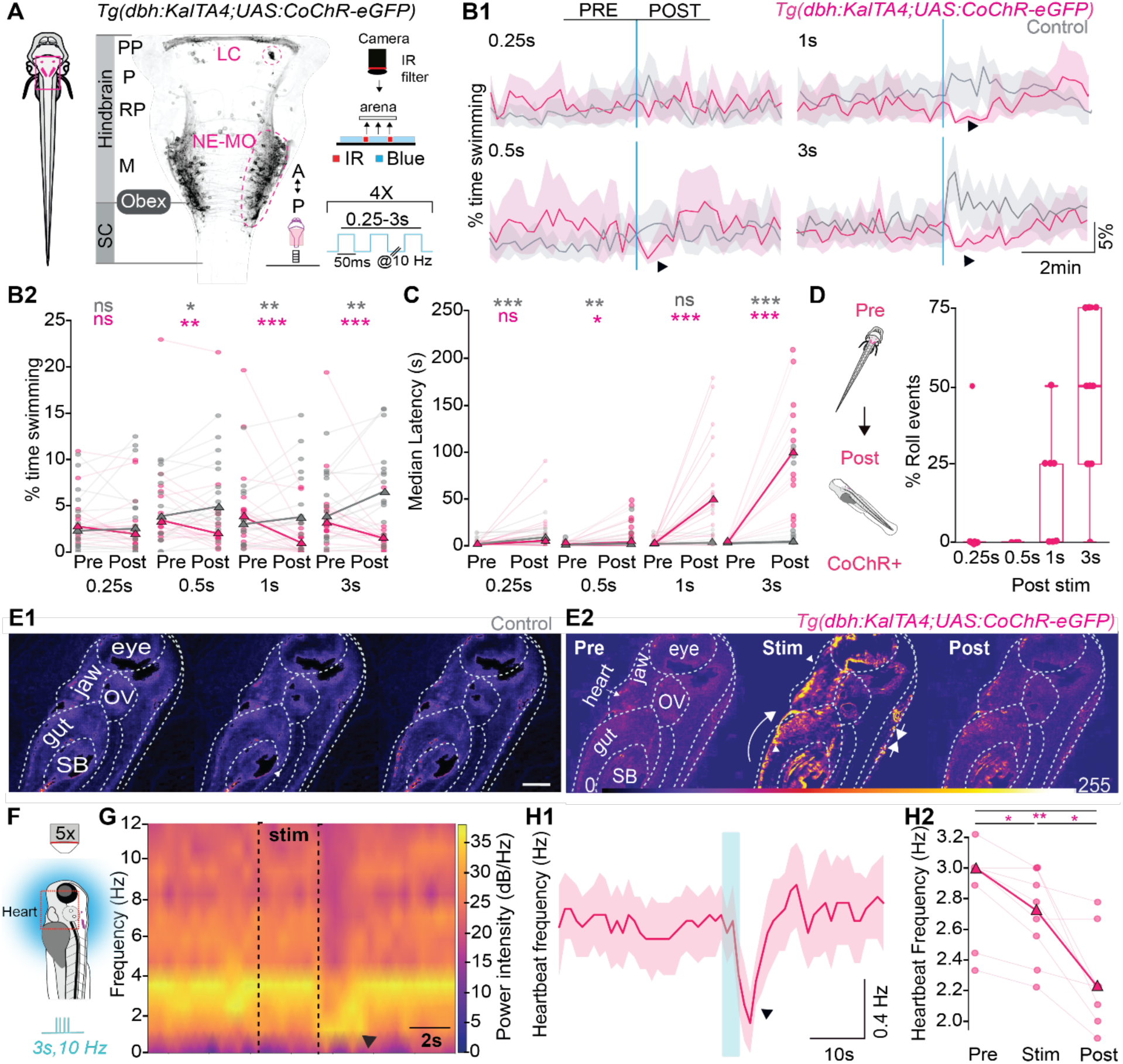
Sustained noradrenergic neuron activation is sufficient to induce a global body response including prolonged motor arrest, loss of posture, emetic reflex and bradycardia. **(A)** Schematic of the experimental paradigm for monitoring locomotor activity altered by optogenetic stimulation of noradrenergic neurons in enucleated 6 dpf *Tg(dbh:KalTA4;UAS:CoChR-eGFP)* zebrafish larvae. Larvae were tracked at 100 Hz in a multi-well arena using infrared light emitting diode (LED) backlight illumination, while blue LEDs (470 nm) delivered intermittent optogenetic stimulation during high-speed video acquisition (see STAR Methods). SC, spinal cord; PP, prepontine; P, pontine; RP, retropontine; M, medulla; A, anterior; P, posterior. Scale bar, 100 μm. **(B1)** Optical train stimulations lasting 0.5 s or more in *Tg(dbh:KalTA4;UAS:CoChR-eGFP)* larvae robustly induced motor arrest, while they increased locomotion in control siblings. Locomotor activity quantified as a fraction of time spent swimming per 10 s bin for varying optogenetic train stimulation durations. Traces show first stimulation (N = 16 control, 16 transgenic larvae; 4 stimulations per larva), see also Figure S1 for individual trial comparisons. **(B2)** Optical train stimulations lasting at least 0.5 s significantly decreased swimming in transgenic larvae but increased swimming in controls: quantification of the fraction of time spent swimming during 120 s-long epochs before and after optogenetic train stimulation (Wilcoxon signed-rank test; 0.25s: control p = 0.28, transgenic p = 0.23; 0.5s: control p = 0.044, transgenic p = 0.013; 1 s: control p = 0.010, transgenic p = 0.00058; 3 s: control p = 0.0042, transgenic p = 0.000061). **(C)** Arrest duration increased in transgenic larvae for optical train stimulations lasting 0.5 s and above. Motor arrest duration (median time from optogenetic stimulus onset to subsequent swim bout) compared to baseline median interbout interval (n = 16 control, 16 transgenic; 4 stimulations per larva; Wilcoxon signed-rank test; 0.25s: control p = 0.00030, transgenic p = 0.052; 0.5s: control p = 0.0026, transgenic p = 0.047; 1 s: control p = 0.060, transgenic p = 0.00021; 3 s: control p = 0.000070, transgenic p = 0.000010). **(D)** Rolling behavior induced by 3 s-long, 10 Hz optogenetic train stimulation in *Tg(dbh:KalTA4;UAS:CoChR-eGFP)* larvae. The cartoon illustrates larval body position before and after light onset. Boxplots show fraction of stimuli inducing rolling for each train duration (n = 16 control, 16 transgenic larvae; 4 stimulations per larva; center line, median; box, 25th–75th percentiles; mean ± SEM: 0.25s, 3.8 ± 3.8%; 0.5s, 0%; 1 s, 9.6 ± 4.5%; 3 s, 44.2 ± 7.0%). **(E1-E2)** Sustained noradrenergic neuron train activation elicits contractions in peripheral organs and musculature. **(E1)** Representative standard deviation of pixel intensities before, during, and after optogenetic train stimulations in control larvae. **(E2)** Representative standard deviation of pixel intensities before, during, and after sustained noradrenergic neuron activation shows contractions in gut, intestine, jaw, heart, and rostral trunk. **(F-G)** 3s-long 10 Hz train optogenetic stimulation also induces bradycardia in transgenic fish. **(F)** Experimental paradigm for monitoring heart rate (see STAR Methods). **(G)** Spectrogram of ROI pixel intensities post Fourier transformation (see STAR Methods) shows a transient reduction in frequency lasting few seconds after sustained optogenetic activation of noradrenergic neurons in *Tg(dbh:KalTA4;UAS:CoChR-eGFP)* larvae (50ms-long pulse at 10 Hz for 3s; n= 4 fish, 8 trials). **(H1)** Sustained activation of noradrenergic neurons reduced the heartbeat frequency of *Tg(dbh:KalTA4;UAS:CoChR-eGFP)* larvae. Frequency of heartbeat computed per time 1s-long bin promptly reduces after the 3s-long optical stimulation. **(H2)** Quantification of mean frequency during 3 s bin pre, during and post optical train stimulation shows a mild reduction in frequency during optical train stimulation which is augmented post train stimulation. Statistical comparisons were made with Wilcoxon signed-rank test (pre vs stim, statistic: 0, p-value: 0.01755, stim vs pre: statistic: 0, p = 0.01755, pre vs stim: statistic:0, p = 0.0078).

### Sustained noradrenergic neuron activation recruits medullary vagal motor neurons but not reticulospinal neurons

To identify the neural circuits mediating these coordinated autonomic responses and induction of motor arrest, we simultaneously performed whole-brain calcium imaging and sustained optogenetic stimulation (10Hz, 3s) in enucleated *Tg(dbh:KalTA4;UAS:CoChR-eGFP;elavl3:jRGECO1b)* larvae at 6 dpf **(Figure 2A)**. Blue light (light emitting diode at 460 nm) absorbed by mApple-based calcium sensors such as jRGECO1b results in an artefactual fluorescence increase upon illumination independent of neuronal activity ^55^. To correct for this artefact, we developed a preprocessing pipeline using principal component analysis (PCA) to isolate and subtract the dominant artefact (first principal component) and used the corrected fluorescence signals for further analysis (see Star Methods, **Figure 2B1–B2, S2A,B**). Control enucleated *Tg(elavl3:jRGECO1b)* siblings showed minimal recruitment (**Figure S2C1**) upon blue light illumination while *Tg(dbh:KalTA4;UAS:CoChR-eGFP;elavl3:jRGECO1b)* larvae showed recruitment of neurons distributed from the optic tectum to the hindbrain but not in the spinal cord (**Figure 2C, Figure S2C2**). Noradrenergic neurons in the medulla oblongata, locus coeruleus, area postrema and cranial nuclei themselves were recruited upon optogenetic activation, serving as a positive control (**Figure S2D**). Anatomical registration with motor neuron -specific transgenic lines revealed that neurons recruited in the caudolateral medulla colocalized with vagal motor neurons (**Figure 2D1-D3; S3A).** Functionally, extracellular electrophysiological recordings confirm sustained noradrenergic activation elicited neuronal activity in visceral motor neurons of the rostral spinal cord segments 6-7 during the stimulation (**Figure 2E1-E3**).Reticulospinal neurons, including the *vsx2⁺* neurons in the brainstem (**Figure S3B)**, provide descending excitatory drive to spinal CPGs for locomotion^35,56,57^. Sustained noradrenergic activation failed to recruit *vsx2*+ reticulospinal neurons eliciting locomotion^35^(**Figure 2F**). Consistently, as observed in freely swimming and enucleated *Tg(dbh:KalTA4;UAS:CoChR-eGFP)* larvae, optogenetic activation of noradrenergic neurons failed to induce tail undulations (**Figure 2G1-G2**). Together, these results demonstrate that sustained noradrenergic activation recruits vagal motor neurons to elicit autonomic outputs but not reticulospinal neurons driving locomotion.

**Figure 2.**
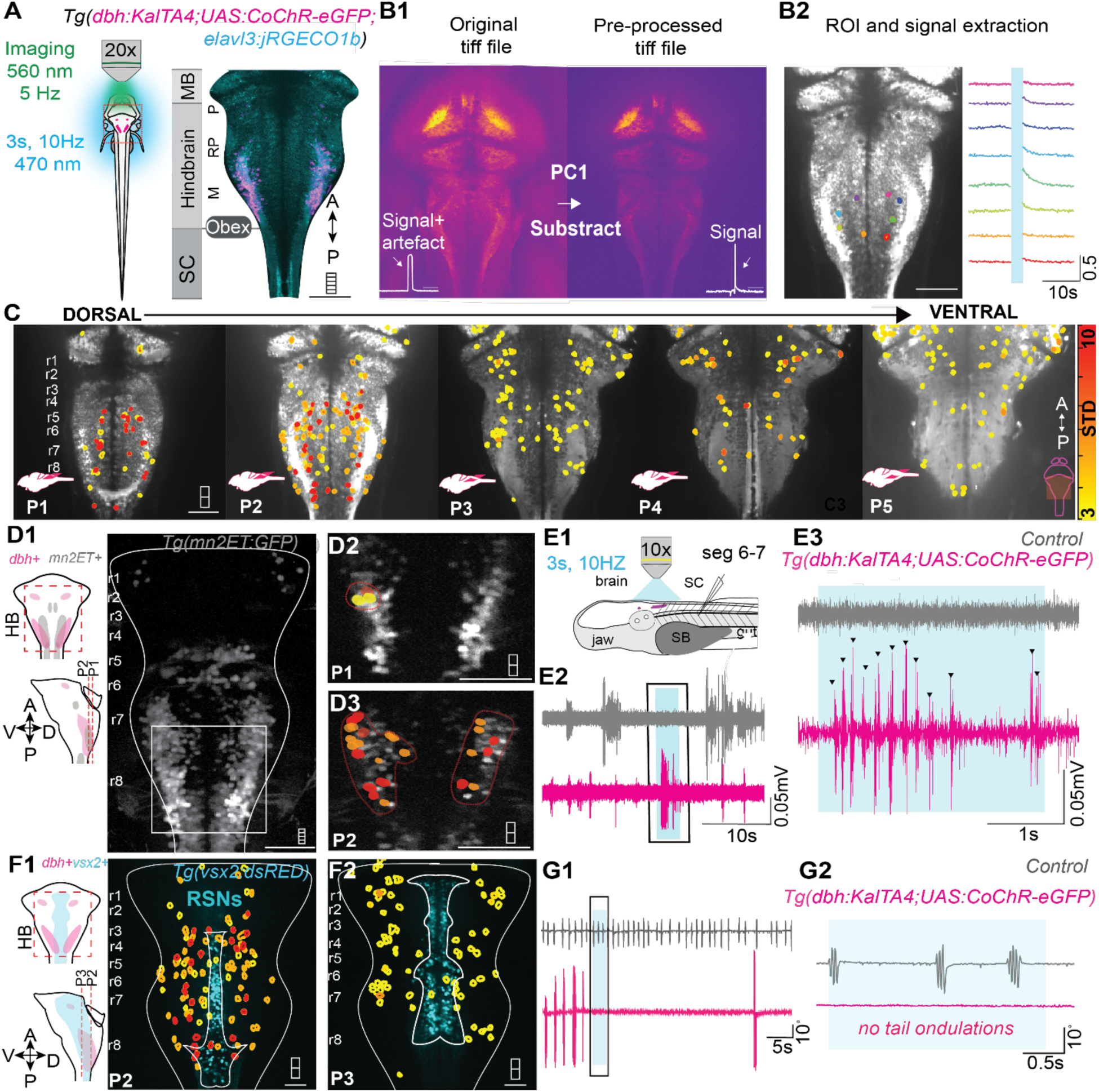
Sustained noradrenergic neuron activation recruits medullary vagal motor neurons but fails to recruit *vsx2^+^* reticulospinal neurons. **(A)** Experimental setup for the simultaneous sustained optogenetic activation of noradrenergic neurons and whole-brain calcium imaging in enucleated *Tg(dbh:KalTA4;UAS:CoChR-eGFP;elavl3:jRGECO1b)* larvae. MB, midbrain; SC, spinal cord; P, pontine; RP, retropontine; M, medulla; A, anterior; P, posterior. Scale bar, 100 μm. **(B1)** Preprocessing step to remove optogenetic artifact due to the blue light absorption of jRGECO1b from the acquisition signal. Subtracting the first principal component PC1 from the original time series yields a clean signal. **(B2)** ROI segmentation and signal extraction from detected ROIs using Suite2p software. Representative calcium traces are shown. Scale bar, 100 μm. **(C)** Sustained activation of noradrenergic neurons shows recruitment of distributed neurons across z-planes in a *Tg(dbh:KalTA4;UAS:CoChR-eGFP;elavl3:jRGECO1b)* larva. Recruitment map color-coded based on response amplitude from baseline. Five imaging planes (P1–P5) covered a depth of 111 µm with an average inter-plane spacing of ∼28 µm. Scale bar, 100 μm. **(D1–D3)** Sustained noradrenergic neuron activation recruits vagal motor neurons in the caudal hindbrain. Subset of recruited neurons in the caudolateral reticular formation juxtaposed with *mn2^ET+^* vagal motor neurons. Scale bars, 100 μm. **(E1–E3)** Sustained noradrenergic neuron activation elicits successive autonomic efferent activity. **(F1)** Experimental setup for extracellular recordings from autonomic efference in rostral spinal cord segments 6–7. **(F2–F3)** Representative extracellular recordings show persistent activity during sustained noradrenergic neuron activation. **(F1-F2)** Sustained noradrenergic neuron activation fails to recruit *vsx2+* reticulospinal neurons in the hindbrain. Minimal overlap of recruitment maps with anatomical location of *vsx2+* neurons. Scale bar, 100 μm. **(H1–H3)** Sustained noradrenergic neuron activation fails to induce tail undulations representative of locomotion in enucleated *Tg(dbh:KalTA4;UAS:CoChR-eGFP)* larvae. **(H1)** Experimental setup for optogenetic activation of noradrenergic neurons with simultaneous monitoring of tail activity. **(H2–H3)** Representative tail undulations in control sibling **(H2)** and *Tg(dbh:KalTA4;UAS:CoChR-eGFP)* larva **(H3)** show no induction of tail undulations upon optogenetic activation (10 Hz, 3 s).

### Motor suppression from sustained noradrenergic activation occurs via reduced synaptic drive on motor neurons

To determine how motor circuits are disrupted upon sustained activation of noradrenergic neurons, we analyzed the modulation of locomotor activity at the level of spinal motor neurons (**Figure 3A**). We first tested whether motor arrest results from suppressed activity in motor neurons. Ventral nerve root recordings from midway along the length of the body (spinal segments 14-15) of paralyzed *Tg(dbh:KalTA4;UAS:CoChR-eGFP)* larvae during optogenetic activation (10 Hz, 3 s) confirmed robust suppression of motor neuron firing in transgenic larvae compared to controls (**Figure 3B-C**). Fraction of swimming time (**Figure 3D1, 3D2**), and motor arrest duration (control 1.776 ± 0.843 s, opsin 21.79 ± 6.83 s; **Figure 3E**) confirmed motor output suppression at the spinal level. To determine whether the silencing of fictive locomotion resulted from the direct silencing of spinal neurons, we performed *in vivo* whole-cell recordings from primary motor neurons in spinal segment 14-15 of *Tg(dbh:KalTA4;UAS:CoChR-eGFP;mn2ET:GFP)* larvae (**Figure 3F**). Input resistance remained unchanged during optogenetic activation (LED OFF: 94.25 ± 2.25 MΩ; LED ON: 94.25 ± 2.25 MΩ; **Figure 3G1-G2**). Spiking elicited by current injections of 300 pA (determined by rheobase measurements)) were consistent between stimulation conditions (**Figure 3H-I**), and analysis of spike properties revealed no significant changes (**Figure 3J-N**), These results demonstrate an absence of indirect modulation of intrinsic properties of the spinal motor neurons. Furthermore, optogenetic stimulation was not associated with mono- or polysynaptic inputs recorded in both primary and secondary spinal motor neurons (data *not shown*), ruling out an inhibitory modulation of motor neurons activity by noradrenergic neurons activation. We next tested whether the sustained noradrenergic activation reduced synaptic inputs to motor neurons by recording motor neurons activity in current clamp configuration. Neurons were held at - 68 mV away from to the reversal potential of chloride (−50 mV in our conditions). Sustained noradrenergic activation (10Hz, 3s), led to a strong reduction of spontaneous post synaptic potentials (PSPs) that persisted for tens of seconds, after an initial likely visual response (as these larvae were not enucleated) (**Figure 3O,Q; Figure S4;** quantified as the mean ± SEM integral of membrane potential : control pre: 18.56 ± 3.05 mV·s, post: 20.16 ± 1.75 mV·s; opsin pre: 22.16 ± 2.47 mV·s, post: 10.13 ± 5.56 mV·s; **Figure 3P1-P2**). Regardless, the suppression of both spontaneous and visually evoked synaptic drive was observed in transgenic larvae compared to control siblings (**Figure 3R1-R2;** control: 22.62 ± 5.28 mV; opsin: 2.09 ± 3.44 mV). Together, these results demonstrate that sustained noradrenergic activation suppresses motor output by reducing the synaptic drive to spinal motor neurons without altering their intrinsic excitability or recruiting synaptically connected inhibitory interneurons.

**Figure 3.**
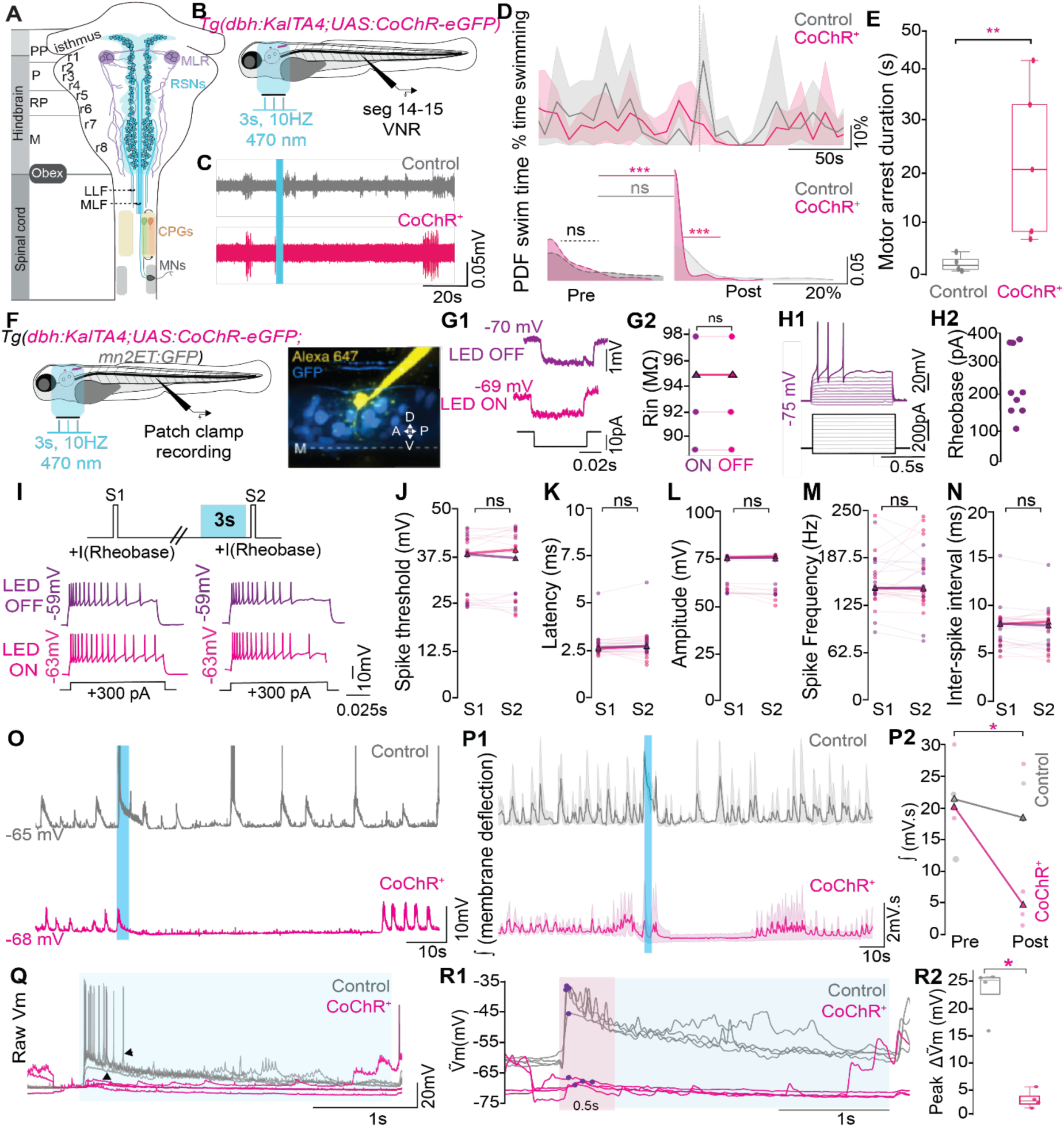
Sustained noradrenergic neuron activation suppresses the synaptic inputs to motor neurons without altering their intrinsic properties. **(A)** Schematic representing the descending motor workflow from brainstem-localized excitatory motor circuits to spinal motor circuits. MLR, mesencephalic locomotor region; RSNs, reticulospinal neurons; CPGs, central pattern generators; MNs, motor neurons; LLF, lateral longitudinal fasciculus; MLF, medial longitudinal fasciculus; PP, prepontine; P, pontine; RP, retropontine; M, medulla. **(B)** Experimental setup showing a ventral nerve root (VNR) recording from segment 14–15 in a paralyzed *Tg(dbh:KalTA4;UAS:CoChR-eGFP)* larva subjected to sustained global activation of noradrenergic neurons. **(C)** Representative VNR traces showing suppression of motor output at segment 14–15 in transgenic larvae compared to control siblings. **(D)** Top panels. Fraction of time spent swimming computed over 10 s intervals demonstrates reduction in swimming upon sustained optogenetic activation of noradrenergic neurons in *Tg(dbh:KalTA4;UAS:CoChR-eGFP)* larvae compared to control siblings. Bottom panels. Locomotor activity computed over 60 s decreases for *Tg(dbh: GAL4;UAS:CoChR-eYFP)* larvae after optogenetic stimulation but not in control siblings (control n=4, opsin n=5, 3 trials per larva). Kolmogorov-Smirnov test, control pre vs post p = 0.087, opsin pre vs post p = 8.06 × 10⁻⁷, control vs opsin pre p =0.093, control vs opsin post p = 6.42 × 10⁻⁵. **(E)** Motor arrest duration, computed as the duration to the first locomotor episode post-optogenetic stimulation, shows a significant increase in *Tg(dbh:KalTA4;UAS:CoChR-eGFP)* larvae compared to control siblings. Mann-Whitney U test: U = 20.0, p = 0.0079 **(F)** Experimental setup for *in vivo* targeted patch-clamp whole-cell recordings from primary motor neurons in spinal cord segment 14 paired with optogenetic activation of noradrenergic neurons (10 Hz, 3 s-long) in *Tg(dbh:KalTA4;UAS:CoChR-eGFP;mn2ET:GFP)* larvae. **(G1–G2)** Sustained optogenetic activation does not change input resistance of spinal motor neurons. (G1) Representative membrane deflection induced by current step injection (10 pA) with and without optogenetic activation of noradrenergic neurons. (G2) Input resistance values remain unchanged with and without optogenetic activation (n = 4 cells, LED OFF: mean = 94.25 ± 2.25 MΩ; LED ON: mean = 94.25 ± 2.25 MΩ. Mann-Whitney U test: p = 1, U statistic =0.0). **(H1–H2)** Typical firing pattern of primary motor neurons upon current injection step of 300 pA (H1) and rheobase of recorded motor neurons (H2). **(I)** Spiking of a typical primary motor neuron elicited by minimum current injection step (rheobase, 300 pA) is consistent over time (top) and does not change after optogenetic train stimulation (10 Hz, 3 s; S1 vs. S2, bottom). **(J–N)** No changes in intrinsic properties of motor neurons upon sustained optogenetic activation of noradrenergic neurons. (J) Spike threshold. (K) Latency. (L) Amplitude. (M) Spike frequency. (N) Inter spike interval. All parameters remain similar after optogenetic activation. Wilcoxon tests p values in Supplemental table 2, n = 3 cells, 3 fish, 12 trials total. **(O)** Representative traces of membrane potential recorded in current-clamp configuration from a primary motor neuron in *Tg(dbh:KalTA4;UAS:CoChR-eGFP;mn2ET:GFP)* larva and control sibling, showing reduction of synaptic inputs to primary motor neurons upon optogenetic activation of noradrenergic neurons lasting tens of seconds. **(P1)** Integral of membrane deflections of all trials and neurons recorded. (P2) Reduction in area under the curve in *Tg(dbh:KalTA4;UAS:CoChR-eGFP;mn2ET:GFP)* larvae but not in control siblings. n = 2 cells, 4 trials. Paired t-test control pre vs post p = 0.6, opsin pre vs post p = 0.033. **(Q)** Zoomed overlaid traces of all trials during optogenetic stimulation show membrane depolarization due to visual response to light in control siblings that is attenuated in *Tg(dbh:KalTA4;UAS:CoChR-eGFP;mn2ET:GFP)* larvae. **(R1)** Membrane potential traces after low-pass filtering to isolate slow depolarization dynamics show a depolarization in the control siblings (visual responses) that is largely reduced in the transgenic larvae. **(R2)** Peak membrane deflection from baseline to 0.5s post-stimulus onset is significantly decreased in *Tg(dbh:KalTA4;UAS:CoChR-eGFP;mn2ET:GFP)* larvae compared to control siblings (Mann-Whitney U, U=16, p = 0.028).

### Sustained noradrenergic activation interrupts descending brainstem motor commands to the spinal cord

To investigate whether noradrenergic neurons suppress motor output by disrupting descending commands from the brain to spinal cord, we established a stimulation method to reliably activate spinal projecting neurons in the brainstem (**Figure 4**). To validate our approach, we paired targeted electrical stimulation of the obex, region at the boundary of hindbrain and spinal cord whose stimulation is effective at initiating locomotion^58^ with the recording of *vsx2^+^* reticulospinal neurons involved in driving the recruitment of spinal motor circuits^35,57,59,60^. We confirmed the recruitment of *vsx2+* reticulospinal neurons by monitoring population activity during electrical stimulation in the obex (2 ms-long pulses at 10 Hz for 40 s) in *Tg(vsx2:GAL4;UAS:GCaMP6s)* larvae (**Figure 4 A, B1, B2)**. We next tested whether sustained noradrenergic activation disrupts electrically evoked descending drive by combining electrical stimulation (2 ms-long pulses at 10 Hz for 40 s) with optogenetic train stimulation (10 Hz for 3 s) in *Tg(dbh:KalTA4;UAS:CoChR-eGFP)* larvae (**Figure 4C**). While stimulation at the obex alone (“LED OFF”) reliably evoked robust tail undulations (**Figure 4D1**), sustained noradrenergic activation (“LED ON”) abolished the motor response (**Figure 4D2**), with that a refractory period during which locomotion could not be evoked (**Figure 4E1).** Noradrenergic stimulation reduced the fraction of time spent swimming (**Figure 4E2**; LED OFF pre mean = 38.70 ± 6.43 %, post mean = 54.18 ± 6.30 %, LED ON pre mean = 40.90 ±8.63 %, post mean = 19.82 ±10.72 %). The refractory period estimated as the time point at which the LED ON and OFF traces intersected lasted 15.6 ± 2.46 s (**Figure 4E3**). These results demonstrate that sustained noradrenergic activation establishes a motor refractory period during which the descending motor commands driven by reticulospinal neurons in the brainstem are disrupted.

**Figure 4.**
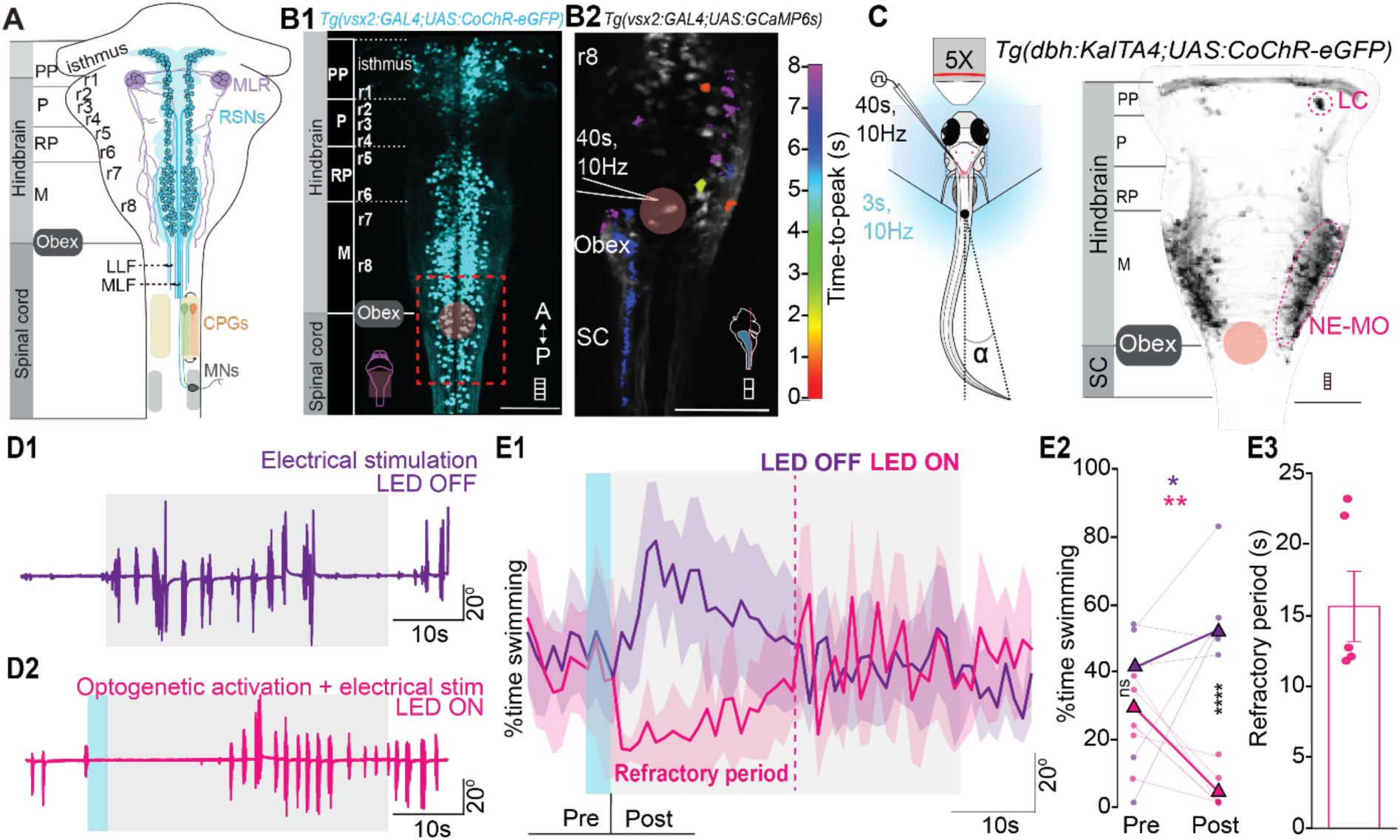
Sustained noradrenergic neuron activation forbids descending motor commands from brainstem to spinal cord. **(A)** Schematic representing the descending motor workflow from brainstem-localized excitatory motor circuits to spinal motor circuits. MLR, mesencephalic locomotor region; RSNs- reticulospinal neurons; CPGs, central pattern generators; MNs, motor neurons; LLF, lateral longitudinal fasciculus; MLF, medial longitudinal fasciculus; PP, prepontine; P, pontine; RP, retropontine; M, medulla. **(B1)** Maximum intensity projection of *Tg(vsx2:GAL4;UAS:CoChR-eGFP)* larva showing the anatomical location of vsx2+ reticulospinal neurons in the caudal brainstem. A, anterior; P, posterior. Scale bar, 100 μm. **(B2)** Control calibration experiment showing induction of descending drive upon electrical stimulation of the obex region (2 ms-long pulse at 10 Hz for 40 s) in *Tg(vsx2:GAL4;UAS:GCaMP6s)* larva. SC, spinal cord. Scale bar, 50 μm. **(C)** Experimental paradigm used to test the effect of sustained noradrenergic activation on induction of motor responses by electrically stimulating the obex region (2 ms-long pulse at 10 Hz for 40 s) in *Tg(dbh:KalTA4;UAS:CoChR-eGFP)* larvae. **(D1–D2)** Representative tail angle traces upon electrical stimulation of the obex region alone (D1) and upon sustained activation of noradrenergic neurons followed by electrical stimulation of the obex region (D2) in *Tg(dbh:KalTA4;UAS:CoChR-eGFP)* larva. **(E1–E3)** Sustained noradrenergic neuron activation induces a refractory period during which locomotion cannot be induced through obex stimulation. **(E1)** Fraction of time spent swimming over 1 s time bins increases upon electrical stimulation alone (“LED OFF”, purple) and decreases upon combination of sustained activation of noradrenergic neurons followed by electrical stimulation of the obex (“LED ON”, pink). **(E2)** Quantification of fraction of time spent swimming 7 s before and after optogenetic stimulation shows an increase while LED OFF and a decrease while LED ON (n= 6 fish, mean fraction ± SEM LED OFF pre = 38.70 ± 6.43 %, post = 54.18 ± 6.30 %, LED ON pre mean = 40.90 ±8.63 %, post mean = 19.82 ±10.72 %; Wilcoxon test, LED OFF pre vs post p = 0.15, Wilcoxon test, LED ON pre vs post p = 0.031. (E3) Refractory period, computed as the time at which the LED OFF and LED ON conditions intersect, lasts 15.6 ± 2.46 s.

### The noradrenaline-induced glial calcium activity persists longest at the obex and ventricular midline

As previously shown^49,51^, noradrenergic-mediated motor suppression occurs together with a slower glial signal induced by noradrenergic activation. In the brain, glial calcium waves modulate synaptic transmission through calcium-dependent gliotransmission ^61,62^.To identify the location where the glial calcium wave was most likely to modulate neuronal activity for extended periods of time, we performed simultaneous optogenetic stimulation of noradrenergic neurons (10 Hz for 3 s) and calcium imaging of glia in enucleated *Tg(dbh:KalTA4;UAS:CoChR-eGFP;gfap:jRGECO1b)* larvae (**Figure 5A**). Optogenetic activation of noradrenergic neurons elicited a robust glial calcium response throughout the hindbrain and rostral spinal cord (**Figure 5B**). To unbiasedly characterize the spatiotemporal organization of the glial calcium wave, we developed a pixel-based k-means clustering analysis of the time series optimized by comparing stability scores across k-values to a noise floor (**Figure S4**). This analysis identified six functional clusters, referred to as C1 to C6, according to time-to-peak, with distinct temporal dynamics and spatial localization (**Figure 5B, C1-2)**. As expected, the earliest responding cluster spatially overlapped with noradrenergic neuron projections in the brainstem and spinal cord (cluster 1, **Figure 5C1-C2**). Spatial maps corresponding to each cluster across all larvae (**Figure 5D1**) revealed that spinal glial activation exhibited the shortest time-to-peak (Clusters C1: 6.28 ± 0.52 s; C2: 7.32 ± 0.49 s, C3: 9.3 ± 0.68 s), while hindbrain populations (C4: 10.96 ± 0.73 s, C5: 12.36 ± 1.09 s, C6: 16.06 ± 1.01 s; **Figure 5D2**) peaked later. To identify the regions with prolonged activity, we color-coded pixels based on their full width at half maximum of the calcium transient. The obex region at the boundary of hindbrain and spinal cord together with the midline in the hindbrain exhibited significantly prolonged calcium signals (obex: 8.89 ± 0.21 s; midline: 12.93 ± 0.37 s) compared to other regions (rostral hindbrain: 5.13 ± 0.098 s; spinal cord: 2.93 ± 0.059 s) (**Figure 5E**; Kolmogorov–Smirnov tests: rostral hindbrain (rHB) vs spinal cord (SC): statistic = 0.4067, p = 8.64 × 10⁻⁷²; rHB vs obex: statistic = 0.3915, p = 1.38 × 10⁻³⁸; rHB vs midline: statistic = 0.7189, p = 4.34 × 10⁻⁵⁷; obex vs SC: statistic = 0.7730, p = 1.03 × 10⁻¹⁴⁴). The sustained calcium activity at the obex and midline approximated the refractory period (15 s) observed during motor suppression (**Figure 4E3**). As expected, short noradrenergic activation for 0.5 s or 1 s failed to sustain glial calcium activity in the obex and midline (**Figure S5A1**; Kolmogorov–Smirnov tests: HB vs SC: statistic = 0.0014, p = 1.0; HB vs obex: statistic = 0.0091, p = 1.0; obex vs SC: statistic = 0.0105, p = 1.0), while 1 s stimulation produced a localized hotspot at the obex but not the midline (**Figure S5A2**; Kolmogorov–Smirnov tests: HB vs SC: statistic = 0.0605, p = 6.9 × 10⁻⁴; HB vs obex: statistic = 0.0982, p = 5.6 × 10⁻⁶; obex vs SC: statistic = 0.1587, p = 2.3 × 10⁻¹⁸). Anatomical overlay of noradrenergic projections showed convergence at the obex (**Figure 5F1, supplementary video 3**), together with targeted projections to the midline (**Figure 5F2, supplementary video 3**). Together, these results uncover that sustained noradrenergic activation induces glial calcium waves that are short-lasting in the spinal cord but persist for tens of seconds in the obex region and midline of the hindbrain.

**Figure 5.**
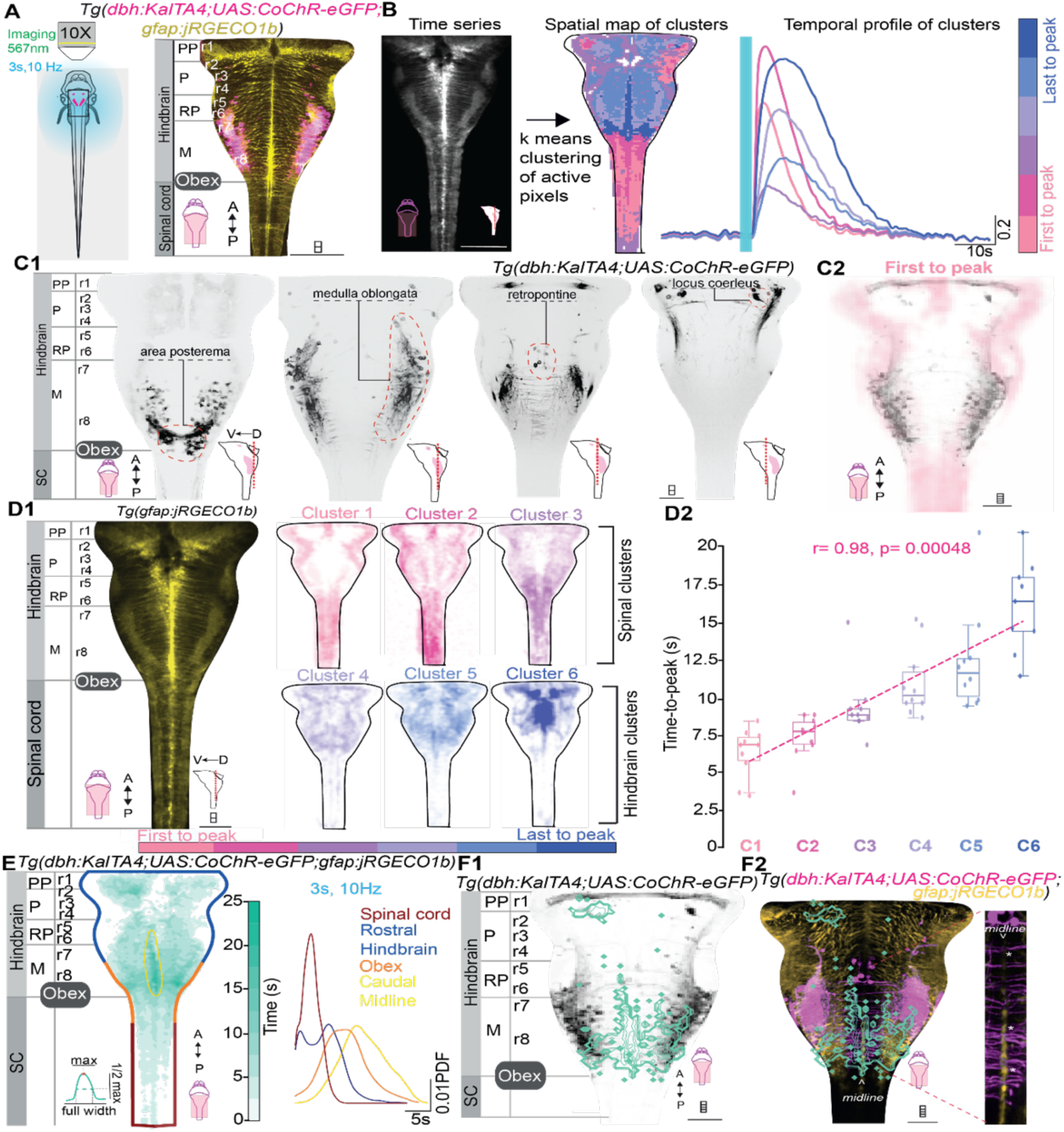
Sustained noradrenergic activation triggers widespread glial calcium waves along noradrenergic projections, lasting in the obex and hindbrain midline. **(A)** Experimental paradigm showing global optogenetic train activation (50 ms-long pulse width, 10 Hz for 3 s) of noradrenergic neurons and simultaneous recording of glial calcium activity in enucleated *Tg(dbh:KalTA4;UAS:CoChR-eGFP;gfap:jRGECO1b)* larvae. Anterior, left; dorsal, top. PP, prepontine; P, pontine; RP, retropontine; M, medulla. **(B)** Pixel-based k-means clustering analysis of time series based on temporal dynamics of calcium signals reveals spatial map of functional clusters (left) with their temporal dynamics (right), color-coded based on time-to-peak (six clusters; see Methods). **(C1)** Single-plane anatomical location of different noradrenergic neuron clusters activated during global optogenetic stimulation (3 s-long train at 10 Hz). Scale bar, 50 μm. **(C2)** Overlap of maximum intensity projection of noradrenergic neuron clusters and projections with Cluster 1, which exhibits the shortest time-to-peak. Scale bar, 50 μm. **(D1)** Spatial map combined from all larvae for each cluster, with higher intensity indicating increased overlap across fish (n = 12 larvae, one stimulation per animal). Spinally localized clusters exhibit shorter time-to-peak, while hindbrain-localized regions exhibit longer time-to-peak. Scale bar, 50 μm **(D2)** Quantification of time-to-peak from all clusters shows that C1–C6 are linearly correlated in time, revealing stereotypic propagation of glial calcium activity. Pearson’s coefficient = 0.98, p = 0.00048. The center line represents the median; box represents the 25th and 75th percentiles. **(E)** Spatial map of sustained glial activity measured as fullwidth at the half maximum of the calcium transient reveals the obex and midline as the loci of the longest lasting glial activity (left) as shown the temporal kinetics of different global anatomical regions (right). Kolmogorov–smirnov tests revealed significant differences between regions rostral hindbrain (rHB) vs spinal cord (SC): statistic = 0.4067, p = 8.64 × 10⁻⁷²; rHB vs obex: statistic = 0.3915, p = 1.38 × 10⁻³⁸; rHB vs midline: statistic = 0.7189, p = 4.34 × 10⁻⁵⁷; obex vs SC: statistic = 0.7730, p = 1.03 × 10⁻¹⁴⁴.A- anterior, P- posterior, D- dorsal, V-ventral. PP-prepontine; P- pontine; RP-retropontine; M-medulla. **(F1-F2)** Overlap with noradrenergic neuron projections with the sustained glial calcium activity in the obex region and noradrenergic neuron projections reaching the midline of the hindbrain. Scale bar, 50 μm.

### Adra1a receptors densely localize on motile cilia at the ventricular midline, where the glial calcium wave persists the most

As the sustained glial calcium wave occurred in proximity to dense noradrenergic innervation, we hypothesized that glial cells express enriched molecular machinery for detecting noradrenaline. α1-adrenergic receptors (Adra1a) couple to Gq/11 proteins, mobilize intracellular calcium via IP3-dependent pathways, and mediate noradrenaline-evoked calcium elevations across different glial subtypes^21,44,58,59^. Hence, we next investigated the distribution of Adra1a using immunohistochemistry in the brainstem (**Figure 6**). Adra1a receptors were enriched in regions with extensive noradrenergic innervation including the medulla oblongata and the obex (**Figure 6A2, A3**), the midline and subventricular zone (**Figure 6A1**), including the choroid plexus (**Figure 6A4, A5; Figure S5B**) and central canal opening (**Figure 6A6**). In the medulla oblongata, careful single-plane analysis confirmed that the Adra1a receptors are distributed in the loci where glial calcium activity lasts the most (**Figure 6B**). Three-dimensional reconstruction revealed pronounced Adra1a receptor enrichment in the ependymal zone surrounding the rhombencephalic ventricle (**Figure 6C1-C3**), consistent with localization to ependymal radial glia (ERG^63^) that extend from the ventricular surface into the parenchyma and have motile cilia in contact with the CSF. Accordingly, the Adra1a receptor in the midline of the obex is localized in cilia of *gfap*+ ERGs in *Tg(gfap:GCaMP6f)* larvae (**Figure 6D1-D3**). Immunostaining against glutamylated tubulin revealed Adra1a colocalization to motile cilia (**Figure 6E1-E3**), indicating that ERGs may detect noradrenergic signals via the CSF. Altogether, our results show that the Adra1a receptors are enriched in the brainstem precisely where the glial calcium wave lasts the most, including the medulla oblongata, the midline, and obex.

**Figure 6.**
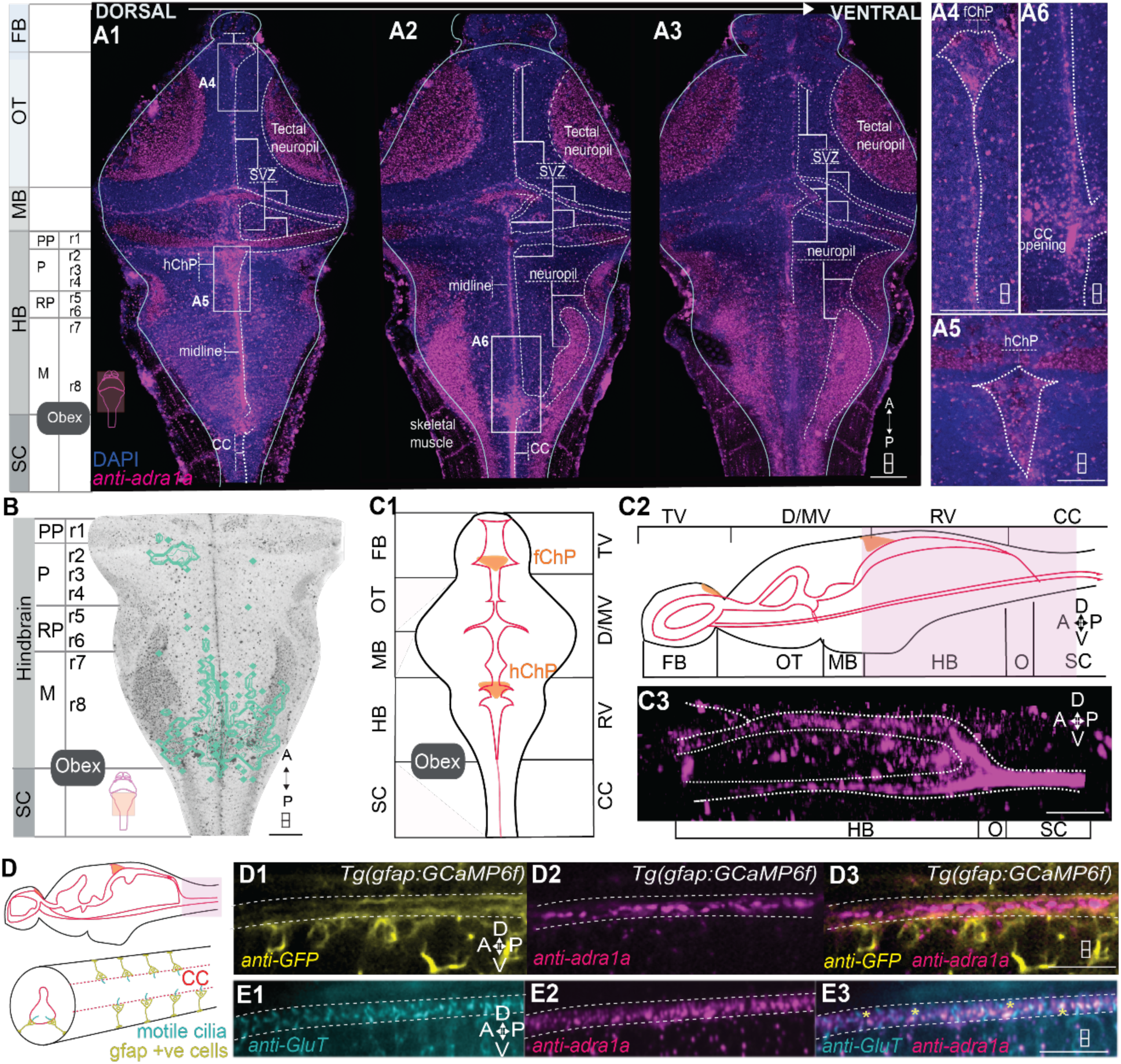
Adra1a receptor distribution overlaps with the sustained glial calcium activity in the obex and midline where it targets motile cilia. **(A1-A6)** Adra1a receptor distribution in the brain and rostral segments of the spinal cord reveals enriched regions in the obex, midline, subventricular zone (SVZ), neuropil and the choroid plexus (ChP). FB, forebrain; OT, optic tectum; MB, midbrain; HB, hindbrain; SC, spinal cord., Scale bar, 50 μm. **(B)** Single plane showing the Adra1a receptor distribution (grey) corresponds to the location of glial calcium activity lasting more than 7s (green). Scale bar, 50 μm. **(C1-C2)** Schematic showing the ventricular system in the brain and central canal in the spinal cord in dorsal (E1) and sagittal view (E2) (TV, telencephalic ventricle; D/MV, diencephalic/mesencephalic ventricle; RV, rhombencephalic ventricle; CC, central canal; A, anterior; P, posterior; D, dorsal; V, ventral) **(C3)** Snapshot of 3D sagittal view of adra1a receptor distribution reveals enrichment of the receptor in the ependymal zone. Scale bar, 50 μm **(D)** Schematic showing the presence of ciliated ependymal radial glia present in the ependymal zone in contact with the cerebrospinal fluid. **(D1-D3)** Single plane projections of the *gfap+ve* cells in the obex showing the localisation of the adra1a receptor in relation to the *gfap+ve* cells in *Tg(gfap:GCaMP6f)* larvae. Scale bar, 50 μm **(E1-E3)** Single plane projections reveal overlap of adra1a with a subset of motile cilia labelled with glutamylated tubulin (anti-GluT). Scale bar, 50 μm

### CSF-mediated noradrenergic signaling prolongs motor arrest

Since noradrenergic neuron activation triggers glial calcium responses localized to Adra1a-expressing ependymal regions, we hypothesized that the cerebrospinal fluid (CSF) could serve for long range signaling of noradrenaline to prolong motor arrest (**Figure 7**). To test whether CSF-borne noradrenaline is sufficient to elicit glial calcium wave, we delivered noradrenaline directly into the rhombencephalic ventricle of *Tg(gfap:GCaMP6f)* larvae. Successful delivery was confirmed by co-injection of Alexa Fluor 547 dextran, which filled the ventricular system without diffusing in the parenchyma (**Figure S6B**). Control injections of artificial CSF (aCSF) failed to evoke glial calcium waves (**Figure S6C1–C2**), confirming response specificity. We next tested a range of noradrenaline concentrations to determine the threshold for CSF-mediated glial activation. Injection of noradrenaline induced concentration-dependent glial calcium responses: low concentrations (1 nM and 5 nM) elicited minimal calcium activity **(Figure S6D1–D2)**. In contrast, injection of 20 nM noradrenaline was sufficient to trigger sustained glial calcium responses, including a long-lasting component in ventricular-contacting region **(Figure 7B; Figure S6D3)**. Time-to-peak analysis revealed that responses initiated at the injection site and propagated along the ventricular system (**Figure 7B1**) with prominent midline hotspots in CSF-contacting regions (**Figure 7B2,** mean ± SEM = 3.064 ± 0.01s), while 430 μM concentrations elicited longer glial calcium activity (**Figure S6 D4; supplementary video 4;** mean ± SEM = 6.55s ± 0.015s) resembling the pattern observed following optogenetic activation of noradrenergic neurons (**Figure 6A**). This spatiotemporal profile corresponds to the Adra1a receptor distribution in the ventricular zone (**Figure 7C)**, suggesting that noradrenaline acts on Adra1a-expressing ERGs.

**Figure 7.**
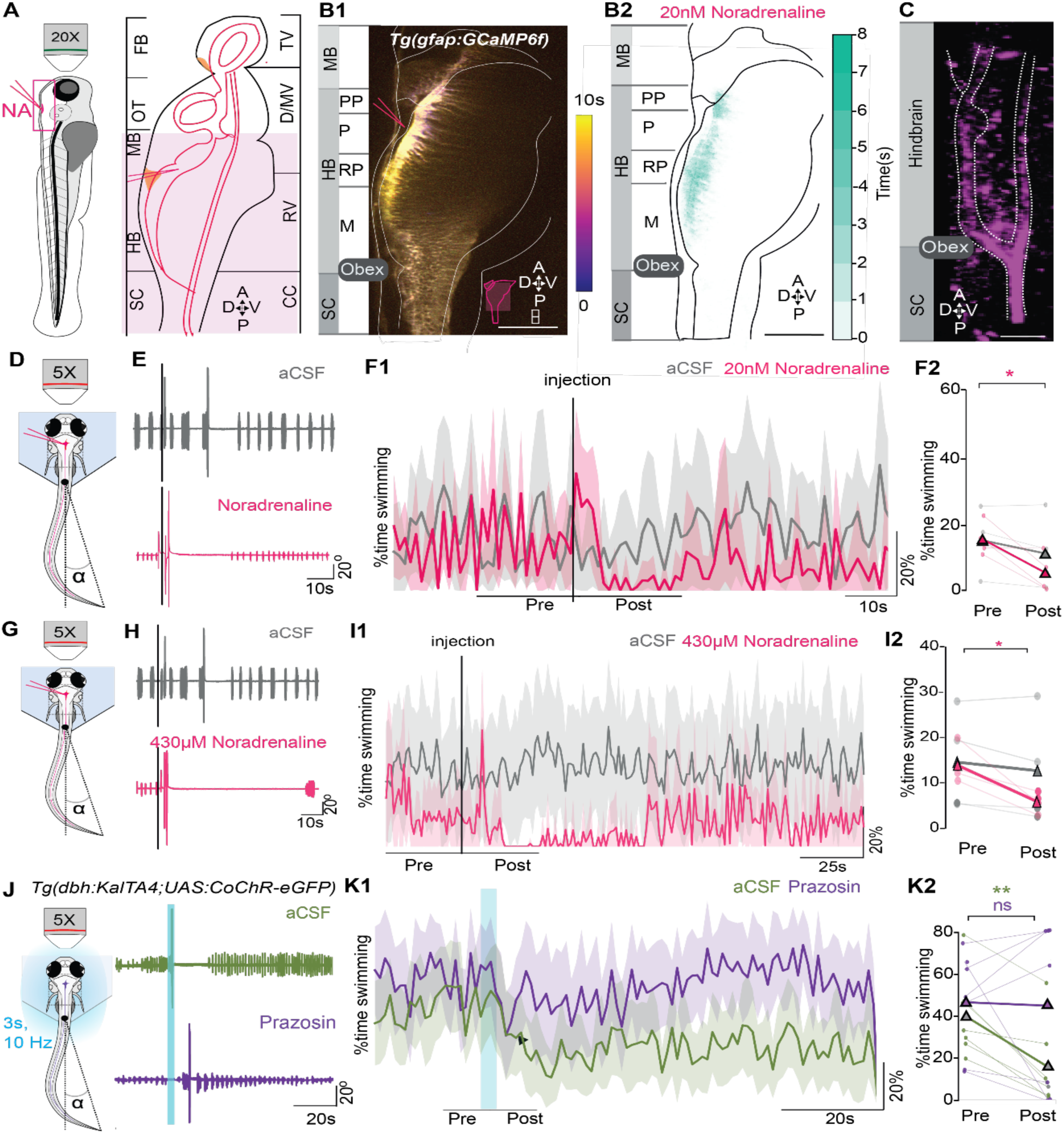
Noradrenaline signaling in the ventricle is sufficient and necessary to prolong motor arrest. **(A)** Experimental paradigm showing the site of ventricular injection of noradrenaline in the rhombencephalic ventricle. FB, forebrain, OT, optic tectum; MB, midbrain;HB, hindbrain; SC, spinal cord; TV, telencephalic ventricle; D/MV, diencephalic/mesencephalic ventricle; RV, rhombencephalic ventricle; CC, central canal; A, anterior; P, posterior; D, dorsal; V, ventral. **(B1-B2)** Ventricular injection of 20 nM noradrenaline elicits glial calcium wave. (B1) Representative time series color-coded based on time-to-peak reveals induction of a widespread glial activity in *Tg(gfap:GCaMP6f)* larvae from the site of injection of noradrenaline (NA) in the rhombencephalic ventricle (n = 4, 2 trials per animal). Scale bar, 100 μm. (B2) Spatio-temporal map of the sustained glial activity reveals hotspots in the midline in contact with the ventricular system. **(C)** The spatio-temporal pattern resembles the adra1a receptor distribution in the ventricular zone. Scale bar, 50 μm. **(D)** Experimental setup to monitor the tail activity upon ventricular injection of noradrenaline. **(E)** Zoomed representative tail angle traces of control aCSF injection (grey) and 20nM noradrenaline injection (pink). **(F1-F2)** Ventricular injection of 20nM noradrenaline evokes motor arrest. (F1) Fraction of time spent swimming over 1 s time bins remains unchanged upon aCSF injection and decreases upon injection of noradrenaline (pink). (F2) Quantification of time spent swimming over 20s epoch before and after stimulation results in a significant decrease in time spent swimming upon ventricular injection of noradrenaline but not aCSF (n = 4, 2 trials per animal, paired t-test, control pre vs post p = 0.078, noradrenaline pre vs post p= 0.035). **(G)** Experimental setup to monitor the tail activity upon ventricular injection of noradrenaline. **(H)** Zoomed representative tail angle traces of control aCSF injection (grey) and 430 μM noradrenaline injection (pink). **(I1-I2)** Ventricular injection of 430 μM noradrenaline evokes motor arrest. (I1) Fraction of time spent swimming over 1 s time bins remain unchanged upon aCSF injection and decreases upon injection of noradrenaline (pink). (I2) Quantification of time spent swimming over 30s epoch before and after stimulation results in a significant decrease in time spent swimming upon ventricular injection of noradrenaline but not aCSF (n=4, 2 trials per animal, paired t-test, control pre vs post p= 0.24, noradrenaline pre vs post p= 0.03). **(J)** Experimental setup to monitor the tail activity upon ventricular injection of prazosin (left), representative tail angle traces of control aCSF injection (green) and prazosin injection (violet). **(K1-K2)** Ventricular injection of 7μM prazosin results in reduction of motor arrest. **(K1)** Fraction of time spent swimming over 1 s time bins induces motor arrest upon aCSF injection and the duration decreases upon injection of prazosin (pink). **(K2)** Quantification of fraction of time spent swimming over 10s epoch before and after stimulation results in a significant decrease in time spent swimming upon ventricular injection of noradrenaline but not aCSF (n = 7, 2 trials per animal, paired t-test, control pre vs post p = 0.001, prazosin pre vs post p = 0.897.

To test whether CSF-delivered noradrenaline was also sufficient to evoke motor arrest, we monitored tail activity following ventricular injection of noradrenaline (**Figure 7D**). Ventricular injection of 0.5nL of 140nM noradrenaline caused motor arrest illustrated by a reduction of tail undulation occurrences compared to aCSF-injected controls (**Figure 7E, F1, F2**; mean ± SEM of fraction time spent swimming in control pre: 15.70 ± 4.65%, control post: 11.81 ± 5.39%, noradrenaline pre: 14.91 ± 2.75%, post: 6.06 ± 2.83%; paired t-test, control pre vs post p= 0.078, noradrenaline pre vs post p= 0.0348), indicating that 20 nM noradrenaline in the CSF is sufficient to evoke motor arrest with higher 430 μM concentration of noradrenaline leading to longer motor suppression (**Figure S6 E-F**; mean ± SEM of fraction time spent swimming in control pre: 14.31 ± 11.14%, control post: 12.61 ± 12.22%, noradrenaline pre: 13.80 ± 4.13%, post: 5.67 ± 2.20%; paired t-test, control pre vs post p= 0.24, noradrenaline pre vs post p= 0.03).

To test whether CSF-mediated noradrenergic signaling is also necessary for long-lasting motor arrest, we injected Adra1 antagonist prazosin to reach a final concentration of ∼7 μM in the rhombencephalic ventricle prior to sustained optogenetic noradrenergic activation (**Figure 7G**). Ventricular prazosin injection significantly reduced motor arrest compared to aCSF-injected controls during sustained noradrenergic activation (**Figure 7H1-2**; mean ± SEM of fraction time in control pre: 40.40 ± 7.33 %,control post: 16.186 ± 19.40%, prazosin pre: 46.771 ± 9.147 %, post: 45.143 ± 15.054 %; paired t-test, control pre vs post p=0.0012, prazosin pre vs post p = 0.8974), indicating that Adra1 receptor activation via the CSF is necessary for the prolonged motor refractory period. Our results establish both sufficiency and necessity of CSF-mediated noradrenergic transmission and support a dual-route model for noradrenaline-mediated disruption of motor circuit function. Noradrenaline acts through sustained glial activity to disrupt descending motor drive within the parenchyma (via synaptic or volume transmission), while simultaneously signaling through the CSF to activate Adra1a in ependymal radial glia. This coordinated action through the parenchyma and CSF ensures a robust, long-lasting motor arrest period following intense threat.

## Discussion

These results establish cerebrospinal fluid as a critical transmission route for coordinating prolonged behavioral states. The sustained glial calcium signaling at the obex, spatially organized by ventricular anatomy and Adra1a receptor localization, suggests a general mechanism by which neuromodulators can maintain circuit-wide suppression without continuous synaptic input. Upon threat, motor arrest consists of coordinated suppression of motor and autonomic systems over long timescales^10,64^ that exceed the typical synaptic transmission timescale^65^. While noradrenergic signaling was known to be involved, how it controlled this prolonged effect was poorly understood^17,66^. We reveal that noradrenaline can orchestrate prolonged motor arrest lasting tens of seconds through two coordinated transmission modes: synaptic signaling in the parenchyma and ciliary signaling via diffusion through the cerebrospinal fluid. The noradrenergic-induced motor arrest is accompanied by postural collapse, bradycardia, and vagal motor neuron recruitment, resembling tonic immobility across vertebrates. Motor arrest does not arise from direct inhibition of motor neurons in the spinal cord but is instead associated with a reduced descending drive from the brainstem to spinal circuits.

Glial calcium activity was the most sustained at the obex and at the midline. Precisely there, along the ventricle walls, Adra1a localizes to the cilia of ependymal radial glial cells, which are present at the interface with the cerebrospinal fluid, a global broadcasting medium for neuromodulatory signals. Accordingly, noradrenaline injected into the hindbrain ventricle was sufficient to recapitulate both the glial calcium wave and motor arrest. Reciprocally, an Adra1 receptor antagonist delivered to the hindbrain ventricle shortened motor arrest upon endogenous noradrenergic activation. Together, these results demonstrate that noradrenaline acts via the parenchyma and CSF to elicit and prolong global motor arrest.

### Sustained noradrenergic activation coordinates a whole-body response through somatic and autonomic suppression

The motor arrest caused by sustained activation of noradrenergic neurons was accompanied by loss of dorsoventral orientation, bradycardia, and emetic-like expulsion behaviors (**Figure 1**). This coordinated suppression of somatic and autonomic output resembles whole-body defensive responses observed across vertebrates during extreme threats ^10^. In humans, severe threat can trigger vagal-mediated collapse with motor paralysis, bradycardia, and nausea^11,26^. Across species, such multi-system responses involve coordinated recruitment of vagal motor outputs alongside sympathoadrenal modulation^12,67^. Interestingly, sustained noradrenergic activation alone, in the absence of any naturalistic threat, is sufficient to recapitulate this multi-system phenotype, revealing that noradrenaline functions as a central coordinator of whole-body defensive states.

Noradrenaline is anatomically positioned to engage this whole-body response through parallel pathways. Centrally, noradrenergic fibers project to vagal motor neurons in rhombomeres 7 and 8^68,69^, which directly innervate the heart and viscera^70^. Peripherally, noradrenergic signaling recruits chromaffin cells in the interrenal gland, the teleost homolog of the adrenal medulla^71,72^, which release catecholamines that suppress cardiac activity via beta-adrenergic signaling^73–75^. This dual architecture enables noradrenaline to coordinate autonomic reconfiguration through both neural and humoral routes. Yet the anatomical site where noradrenaline simultaneously orchestrates loss of postural tone, motor arrest, emesis, and bradycardia has been unclear.

The area postrema at the obex emerges as a strong candidate for this integrative function. This medullary region is critical for both autonomic control and postural maintenance: in mammals, reticulospinal neurons in the caudal medulla provide tonic excitatory drive to spinal motor circuits that sustain muscle tone and posture^76^. The area postrema functions as the primary emetic center, projecting directly to vagal motor nuclei ^77–80^, and AP lesions abolish both emesis and stress-induced cardiovascular changes^81^. The zebrafish obex exhibits analogous connectivity and is activated during our optogenetic stimulation, positioning it to couple motor and autonomic suppression. We propose that noradrenaline acts at this circumventricular site through parallel mechanisms: CSF-mediated activation of Adra1a-expressing ependymal glia maintains the disruption of descending motor commands from medullary reticulospinal centers, leading to loss of postural tone and motor arrest, while direct recruitment of area postrema neurons coordinates vagal outputs driving emesis and contributing to bradycardia. Locus-specific manipulations, such as silencing area postrema neurons while preserving glial Adra1a signaling or blocking glial responses while maintaining area postrema activity, will determine whether motor arrest and autonomic suppression require coordinated action at this site or can be independently controlled.

### Noradrenergic signaling silences motor commands from brain to spinal cord

As shown previously, noradrenergic descending fibers extend longitudinally through ventral and ventrolateral regions of the spinal cord across essentially all spinal segments and are therefore positioned to influence spinal motor circuitry^21,82,83^. To understand how motor output was silenced during motor arrest, we recorded motor neurons via whole-cell patch clamp recordings. We demonstrated that the intrinsic excitability of spinal motor neurons is intact while synaptic inputs are strongly reduced, indicating that motor suppression occurs upstream of the spinal motor pool. This phenomenon could be due to silencing of excitatory inputs to motor neurons or to activation of premotor inhibitory neurons in the spinal cord are recruited. Our data does not support the latter. First, we found no evidence for recruitment of inhibitory interneurons in the spinal cord (**Figure S2 C2**). Second, the glial calcium wave in the spinal cord is short lasting, arguing against the spinal cord as the primary site of sustained suppression. Although our recording conditions cannot formally exclude direct motor neuron inhibition, the lack of change in intrinsic excitability and input resistance suggests that any such effect is likely to have a minor contribution.

Instead, our data supports an alternative explanation: communication between the brainstem and spinal cord is suppressed. We found evidence for long lasting glial calcium wave and high density of Adra1 receptors at the obex, where the axons of reticulospinal neurons project to reach the spinal cord. Stimulation of the obex region is sufficient for motor initiation in adult zebrafish^58^. Accordingly, "start" neurons are present in caudal rhombomere 8 ^35,60^. Electrical stimulation at the obex during noradrenergic activation failed to elicit tail undulations, indicating that noradrenaline interrupts communication between supraspinal motor centers and spinal motor circuits.

Two non-mutually exclusive mechanisms may account for this disruption. First, noradrenergic signaling could recruit GABAergic neurons in the lateral medullary oblongata (L-MO) to inhibit reticulospinal neuron somata, as previously proposed^49,51^. In addition, sustained glial signaling centered at the obex may alter extracellular potassium homeostasis^84,85^ or engage adenosine receptor–mediated presynaptic inhibition^86–88^, thereby suppressing synaptic transmission from descending motor pathways. Regardless of the precise upstream mechanism, the convergence of preserved motor neuron excitability, reduced synaptic drive, and sustained glial calcium activity at the obex support a model in which noradrenaline-mediated glial signaling at the obex enforces motor arrest by selectively disrupting descending motor output.

A recent study in larval zebrafish identified neurons in the ventral prepontine nucleus (vPPN) that regulate tonic immobility duration following repeated vibrational threat^25^. These vPPN neurons express corticotropin-releasing hormone (CRH), a neuropeptide that potently activates locus coeruleus noradrenergic neurons in mammals^89,90^. Our pan-neuronal calcium imaging during sustained optogenetic noradrenergic activation revealed partial vPPN recruitment in one imaging plane (plane 3, **Figure 2**) but not in others. This heterogeneous pattern, together with the finding that vPPN engages specifically during vibrational threat, not electric shock^25^, suggests vPPN functions as sensory modality-specific upstream circuitry rather than a uniform downstream target of noradrenergic signaling. The limited vPPN recruitment during optogenetic activation likely reflects the absence of vibrational sensory input that normally engages this population. This dissociation indicates that while noradrenergic signaling is sufficient to elicit whole-body defensive responses, full recruitment of stress-responsive CRH circuits requires specific sensory features. Whether vPPN provides sensory-gated modulation of noradrenergic tone or operates in parallel to regulate immobility duration independently will be the focus of future studies.

### How does CSF noradrenaline sustain motor arrest beyond synaptic transmission?

Noradrenaline has a half-life of 2.5 minutes in plasma^91^ and slower clearance in CSF, where it lasts hours ^92–94^. Noradrenaline is present in the CSF: it is elevated during stress and in stress-related disorders, including Post-Traumatic Stress Disorder^95,96^. These observations indicate that noradrenergic signaling can extend beyond synaptic transmission into the CSF^65,97^. We demonstrate here that noradrenaline in the CSF is sufficient to induce sustained motor arrest and evoke glial calcium responses. Reciprocally, antagonizing Adra1 receptors shortens motor suppression. Accordingly, we found Adra1a receptors on the cilia of ependymal radial glial, where they can detect ventricular noradrenaline and trigger intracellular IP3-dependent calcium signaling ^61,98^. If noradrenaline signals through the CSF, could noradrenaline also contribute to the long-lasting glial calcium wave in the obex? Adra1a-expressing ependymal glial cells with ciliated apical processes line ventricular walls throughout the brain and are positioned to detect noradrenaline released into CSF, including direct release from noradrenergic neurons in circumventricular structures such as the area postrema^81,99^. While these CSF-contacting ependymal cells are distributed along the entire ventricular system, including the fourth ventricle and central canal, the obex exhibits distinct anatomical features that may concentrate and prolong noradrenergic signaling. The obex constitutes a unique junction where the fourth ventricle narrows into the central canal^100^, spatially concentrating CSF-borne signals^100^. At this transition zone, the Reissner fiber that emerges from the subcommissural organ and extends through the central canal can bind and trap monoamines, including noradrenaline ^101–104^. This could lead to further entrapment of noradrenaline locally at the obex, prolonging its availability to CSF-contacting ependymal cells. The convergence of CSF noradrenaline concentration, monoamine trapping by the Reissner fiber, and the presence of Adra1a-expressing ependymal cells at the ventricular–central canal junction provides a parsimonious explanation for the sustained glial calcium signaling we observe at this site. Critically, the obex is also uniquely positioned where ependymal glial responses to concentrated CSF noradrenaline interface with adjacent reticulospinal neurons that provide descending motor drive.

### Limitations of the study

To identify noradrenergic contribution without sensory stimulation, we employed synchronous optogenetic activation of noradrenergic neurons, an approach that might not recapitulate the spatially and temporally heterogeneous recruitment patterns likely present during naturalistic behavior. Although, ventricular noradrenaline injection experiments were designed to test the sufficiency of CSF-facing adrenergic signaling, endogenous CSF noradrenaline dynamics during noradrenaline-mediated glial calcium wave in larval zebrafish remains to be estimated.

Together, our study shows that noradrenaline functions as a neuromodulator capable of broadcasting a persistent motor suppression signal via the cerebrospinal fluid to sustain ependymal glial calcium activity. This raises a fundamental question: do other monoamines and neuropeptides exploit similar CSF-mediated mechanisms to coordinate brain-wide glial responses? Answering this will be crucial for understanding how the brain integrates whole-body adaptations to threat and stress.

## STAR Methods

### Animal care

The Paris Brain Institute (ICM) and the French National Ethics Committee approved the animal handling and experimental procedures, ensuring compliance with European Union regulations. The approval number for this study was APAFIS #2018071217081175. Adult zebrafish were kept in tanks with a maximum of eight fish per liter. Their environment maintained a temperature of 28.5°C and followed a cycle of 14 hours of light and 10 hours of darkness. Larval zebrafish were typically raised in Petri dishes filled with E3 solution (29.88 mM NaCl, 10.74 mM KCl, 14.57 mM CaCl₂·2H₂O, and 24.05 mM MgCl₂·6H₂O, pH 7.2), under identical temperature and lighting conditions as the adults. The experiments were conducted at 20°C, using zebrafish larvae between 3- and 6-days post-fertilization (dpf). Specific details for each experimental protocol are provided in their respective sections below.

| Transgenic line | Reference |
| --- | --- |
| *Tg(dbh:KalTA4)<sup>jf12Tg</sup>* | Mu *et al.*, 2019<sup>49</sup> |
| *Tg(UAS:CoChR-eGFP)<sup>jf44Tg</sup>* | Mu *et al.*, 2019<sup>49</sup> |
| *Tg(gfap:jRGECO1b)<sup>jf33Tg</sup>* | Mu *et al.*, 2019<sup>49</sup> |
| *Tg(parga<sup>mn2Et/+</sup>:GFP)* | Balciunas *et al.*, 2004<sup>105</sup> |
| *Tg(elavl3:jRGECO1b)<sup>jf17Tg</sup>* | Mu *et al.*, 2019<sup>49</sup> |
| *Tg(vsx2:dsRED)<sup>nns3Tg</sup>* | Kimura *et al.*, 2006<sup>106</sup> |
| *TgBAC(vsx2:GAL4FF)<sup>nns18Tg</sup>* | Kimura *et al.* 2013<sup>59</sup> |
| *Tg(UAS:GCaMP6s)<sup>mpn101Tg</sup>* | Thiele *et al.*, 2014<sup>107</sup> |
| *Tg(gfap:GCaMP6f)<sup>jf22Tg</sup>* | Mu *et al.*, 2019<sup>49</sup> |

#### ELECTROPHYSIOLOGICAL RECORDINGS

##### Loose-patch recordings of CoChR-expressing neurons for calibration of optogenetic stimulation

6 dpf *Tg(dbh:KalTA4;UAS:CoChR-eGFP)* larvae were anaesthetized using MS-222 (MSS-222; MS-222, Sigma-Aldrich) and screened for fluorescent marker expression under a dissection microscope coupled to a LED. Larvae were mounted upright in 3% agarose and paralyzed by muscle injection of α-bungarotoxin (Tocris, 2133). A small window was cut over the brain to permit electrode access. Animals were bathed in artificial cerebrospinal fluid (aCSF; 134 mM NaCl, 2.9 mM KCl, 1.2 mM MgCl₂, 10 mM HEPES, 10 mM glucose, 2.1 mM CaCl₂; 290 mOsm; pH 7.7–7.8 adjusted with NaOH). Skin and dura were dissected using a fine tungsten pin. Patch electrodes were filled with internal solution containing (in mM): 115 K-gluconate, 15 KCl, 2 MgCl₂, 4 Mg-ATP, 10 HEPES free acid, 0.5 EGTA (290 mOsm; pH 7.2 adjusted with KOH), supplemented with Alexa 647 (3 μM final). Fluorescence-guided patch-clamp recordings were performed using an epifluorescence microscope (Nikon, Eclipse FN1). Opsin-positive cells in the medulla oblongata were targeted, and membrane deflections were monitored in a loose-patch configuration. Optogenetic stimulation was delivered as trains of 470 nm illumination with 50 ms pulse width at 10 Hz for durations of 250 ms, 500 ms, 1 s, or 3 s (Prizmatix, UHP-T-HCRI_DI) delivered through the objective.

##### Ventral nerve root recordings

5–6 dpf larvae were paralyzed by bath application of α-bungarotoxin (Tocris) and mounted in 1.5% low-melting agarose. Agar-mounted larvae were bathed in E3 and a portion of agar overlying the spinal cord was removed to allow pipette access. Extracellular ventral root recordings were performed through the skin between myotomes 14 and 18 as described (Masino and Fetcho, 2005). VR pipettes (1B150F-4, WPI) were pulled to record the signals. Signals were digitized at 5-10 kHz using a Multiclamp 700B and a Digidata 1440A (Molecular Devices) and acquired using pClamp10. Three optogenetic stimulations (50ms pulse width, 10Hz, 3s) were delivered per recording in control and transgenic larvae.

###### Analysis

Sweeps (183 s each) were concatenated for subsequent analysis. VR signals were detected as positive peaks exceeding 0.02 (threshold derived from trace noise). Swim bouts were defined as spike clusters with inter spike interval (ISI) ≤ 200 ms and ≥ 2 spikes per bout. Bout duration, bursts per bout, instantaneous burst frequencies, and inter-bout interval (IBI) were computed using custom Python scripts. Stimulus-response latency was computed as time from stimulus offset to onset of the first subsequent bout. Time spent swimming was computed in 1s bins as (sum of bout durations / 1 s) × 100. Pre–post values were computed over 60 s before and 60 s after stimulus onset.

###### Statistical test

Kolmogrov-Smirnov test was used to compare the pre and post time spent swimming and Mann-Whitney U tests were used for arrest duration.

##### *In vivo* whole-cell patch clamp recordings

Whole-cell recordings were performed in 5–6 dpf larvae paralyzed with α-bungarotoxin and pinned in Sylgard-coated, glass-bottom dishes filled with ACSF using thin tungsten pins through the notochord. Skin was removed from segments 5–6 to the end of the tail using fine forceps, and one to two segments were dissected using glass suction pipettes. Patch pipettes (1B150F-4, WPI) were pulled to reach a tip resistance of 5–15 MΩ and filled with internal solution (in mM: 115 K-gluconate, 15 KCl, 2 MgCl₂, 4 Mg-ATP, 10 HEPES free acid, 0.5 EGTA; 290 mOsm; pH 7.2 with KOH) supplemented with 4 mM Alexa 647. Rheobase was measured experimentally by injecting increasing steps of 10pA for 500ms until cells fired action potentials. Rheobase currents were then delivered at two time points (S1 and S2) separated by 1 min. A subset of trials included optogenetic stimulations (460 nm, 3 s) 150ms before the S2 current injection step. Input resistance was measured by injecting a step of −10pA 150ms after the end of the optogenetic stimulation. Current clamp recordings were performed to record spontaneous synaptic activity and synaptic inputs triggered by light emitting diode (LED) stimulations in spinal motor neurons held at −65mV.

###### Analysis

Spike count, spike frequency, median inter spike interval (ISI), median spike amplitude (baseline at pulse onset to peak), and latency to first spike were analyzed using custom Python scripts (pyabf, NumPy, SciPy) and were computed separately for Step 1 (between the first and second pulses) and Step 2 (after the second pulse). Integral of the membrane potential were calculated as the area under the curve using an existing python library and the area under the curve 30s before and after the stimulation was taken. To calculate peak deflection, a low-pass filter was used to the membrane voltage signal and the peaks were detected using Sci-Py detect peaks within 0.5s of the onset of optogenetic stimulation and the peaks were detected and the peak membrane voltage deflection was computed based on the baseline of 0.2s before the onset of stimulation.

###### Statistical test

Wilcoxon paired tests were used for pre-post analysis and Mann-Whitney U test was performed to compute the statistical significance of the difference between opsin and control groups.

#### BEHAVIORAL RECORDINGS

##### Optogenetic stimulation of noradrenergic neurons in freely moving zebrafish larvae

At 3 dpf, larvae were mounted in 3% agarose. Eyes were removed by severing the optic nerve with a custom-made tungsten hook. Animals were bathed in Ringer’s solution (116 mM NaCl, 2.9 mM KCl, 1.8 mM CaCl₂·dH₂O, 5 mM HEPES; pH 7.2) during dissection and allowed to recover for 1 h in Ringer’s solution before transfer to E3. Larvae recovered to 6 dpf, when behavioral recordings were conducted. At 6 dpf, opsin-positive *Tg(dbh:KalTA4;UAS:CoChR-eGFP)* larvae and control siblings were placed in a custom-made 24-well plate (one larva per well in 1 mL E3 to prevent dome formation). The plate was placed in a custom-built behavior setup with constant infrared (IR) background illumination (890 nm; ILH-IW01-85SL-SC211-WIR200, Intelligent LED Solutions) and intermittent blue light (470 nm; ILR-OW16-BLUE-SC211-WIR200, Intelligent LED Solutions). Behavior was recorded at 100 fps using a high-speed camera (acA640-750um, Basler) controlled by Hiris software (RD Vision; https://www.rd-vision.com/r-d-vision-eng). A 600 nm high-pass filter was used to ensure that blue stimulation light did not contaminate imaging. Blue light train stimulations (10 Hz, 50 ms pulse width; durations 250 ms, 500 ms, 1 s, 3 s) were triggered using Arduino control code. Larvae acclimated for 30 min prior to acquisition. Each recording comprised 4 min baseline followed by stimulation at four time points separated by 4-min intertrial intervals (total 20 min). Recordings were repeated with a 20-min delay between acquisitions.

##### Behavioral recordings during electrical stimulations

Larvae were mounted in 3% agarose with tail freed were illuminated laterally at 45° using an 890 nm LED (ILH-IW01-85SL-SC211-WIR200, Intelligent LED Solutions). Behavior was recorded at 300 Hz from above through a 5× objective (5×20/0.25, 12.5-mm; 440125-0000-000, ZEISS) onto an IDS camera (iDS-UI-3060CP-M-GL R2). A 600 nm high-pass filter blocked the 460 nm stimulation LED. Acquisition used Hiris software (RD Vision, St-Maur-Les-Fossés, France, https://rd-vision.com/).

##### Behavioral recordings during ventricle injections

*Tg(gfap:GCaMP6f)* larvae were mounted dorsally, while 150 ms pressure injection of different concentrations of noradrenaline was delivered into the ventricular space, combined with tail tracking. Continuous infrared (IR) backlight illumination (890 nm) was used. Tail undulations were recorded using an IDS camera (iDS-UI-3060CP-M-GL R2) at 100 fps for 180 s using the Hiris software (RD Vision, St-Maur-Les-Fossés, France, https://rd-vision.com/).

Following ventricular prazosin injection, whole field optogenetic stimulation was delivered via a 460 nm LED through the condenser (50 ms - long pulse at 10 Hz for 3 s), triggered by Digidata 1440A controlled by Clampex v10.3. Continuous IR backlight illumination (890 nm) was used. Tail ondulations were recorded using an IDS camera (iDS-UI-3060CP-M-GL R2) at 100 fps for 60 s using the Hiris software (RD Vision, St-Maur-Les-Fossés, France, https://rd-vision.com/). A 600 nm long-pass filter was placed in the detection path to prevent blue-light transmission.

Tail tracking and bout segmentation: Tracking and Analysis: Behavioral videos were tracked using ZebraZoom^108^ (https://github.com/oliviermirat/ZebraZoom). Tracking quality was validated visually. Post-tracking outputs were processed using custom Python scripts.

###### Analysis

Free-swimming optogenetics: Swimming time was computed in 10-s bins as the sum of bout durations per bin divided by bin duration × 100. For pre–post analysis, swimming was quantified over 120 s before and 120 s after stimulation; each larva contributed four pre and four post values, and the median across trials was used per larva. Arrest duration was defined as the time from stimulus onset to the frame of the next swim bout onset. For pre–post analyses, the median IBI during baseline (first 240 s) and the median arrest duration across four trials were used. Rolling events following 3s stimulation were manually flagged. Percent rolling was computed as (rolling events / total events) × 100.

Head embedded behavioral tracking quantification: Time spent swimming was computed in consecutive 1-s bins as (sum of bout durations within bin / 1 s) × 100. Time series plots display mean ± 95% CI across larvae. For pre–post analyses, swimming activity was quantified over the defined period immediately before and after stimulation. Refractory period was computed as the duration at which the electrical stimulation trace intersected with the trace where the electrical stimulation preceded with optogenetic stimulation.

###### Statistical analysis

Paired t-test or Wilcoxon statistical test was performed to compute the statistical significance.

#### CALCIUM IMAGING

##### Neuronal recruitment and glial calcium wave acquisition

Animals were enucleated as described above, immobilized in 3% low-melting-point agarose, bathed in E3, and paralyzed by muscle injection of α-bungarotoxin (Tocris, Bristol, UK). Imaging was performed on a spinning disk confocal microscope (3i, Denver, USA, https://www.intelligent-imaging.com/). TIFF time series were acquired at 5 Hz using a 10× water-immersion objective. Calcium activity was recorded using a 561 nm laser. Whole field optogenetic stimulation was delivered using an intermittent blue LED (466 nm) through the condenser. Light trains (50 ms long pulse at 10 Hz; for 250 ms, 500 ms, 1 s, or 3 s) were generated via digitizer (Digidata 1440A) controlled by Clampex v10.3.

##### Calcium imaging during electrical stimulations

Experiments used an upright microscope (Examiner Z1, ZEISS) with spinning disk head (CSU-X1, Yokogawa, Musashino, Tokyo, Japan) and laser light stack (LaserStack, 3i). Calcium Imaging of *Tg(vsx2:GAL4;UAS:GCaMP6s)* was acquired by using a 20X objective at 5 Hz using 466nm laser.

##### Calcium imaging during ventricle injections

*Tg(gfap:GCaMP6f)* larvae were mounted laterally and imaged on a spinning disk microscope while 150 ms pressure injection of noradrenaline was delivered into the ventricular space, combined with glial calcium imaging. Images were acquired by acquired by a 20X objective at 5 Hz using 466nm laser.

###### Analysis

Quantification of neuronal recruitment: ROIs were detected using Suite2p^109^.To identify neurons recruited by noradrenergic stimulation, we computed a recruitment score (S) for each ROI as the difference between average ΔF/F in the 1-second post-stimulation period and the 1-second pre-stimulation period: S = ⟨ΔF/F⟩{post} - ⟨ΔF/F⟩{pre}. Each experiment was repeated three times per imaging plane. A neuron was classified as recruited if its S score exceeded three standard deviations of the pre-stimulation ΔF/F (calculated over 5 seconds preceding stimulation). To ensure reproducibility, we retained only neurons that met this threshold in all three repetitions, with the final S score computed as the average across repetitions. To anatomically map recruited neurons (imaged in *Tg(elavl3:jRGECO1b)* larvae via spinning disk confocal microscopy) onto defined neuronal populations, we performed sequential correlation-based registration using noradrenergic neuron landmarks. The experimental imaging plane contained both elavl3+ (pan-neuronal) and *dbh+* (noradrenergic) signals. We first registered the *dbh* channel from this plane to a reference z-stack containing *Tg(dbh:KalTA4;UAS:CoChR-eGFP)* and *Tg(vsx2:dsRED*) by computing cross-correlation coefficients across all reference planes and selecting the plane yielding maximum correlation. This registration identified the corresponding vsx2+ anatomical pattern in the matched reference plane. We then registered this *vsx2* pattern to a second reference z-stack containing *Tg(vsx2:dsRED*) and *Tg(mn2ET:GFP),* using the same correlation-based matching approach. The spatial transformation derived from this second registration was applied to identify *mn2ET+* neurons in the coordinate frame of the experimental imaging plane. This sequential registration strategy enabled anatomical mapping of recruited neurons relative to defined neuronal populations.

###### Glial wave calcium imaging analysis

k-means clustering: A custom analysis pipeline was developed to quantify spatiotemporal dynamics of the glial calcium wave using pixel-based k-means clustering (workflow summarized in **Figure S4A**).

###### Pre-processing

Raw TIFF files were spatially binned 4× in each dimension. A larva-encompassing ROI was manually drawn to exclude background pixels. Image series were registered to an internal reference brain using ITK-SNAP^110^ (www.itksnap.org). Optical artifacts during stimulation were addressed by detecting saturated frames, replacing them with the mean of the 5 frames immediately preceding stimulation onset, and treating replaced frames as NaN for subsequent analyses. Motion correction was performed using rigid registration from Suite2p^109^ (v0.8.0; https://github.com/MouseLand/suite2p).

###### k-means clustering

For each file, 400 frames were analyzed (100 pre-stimulus and 300 post-stimulus frames). Background noise was reduced using spatial Gaussian filtering. Activity-based thresholding was used to identify pixels showing activity in the time series. To choose k, stability scores were computed for k = 2–19 across 20 bootstrap runs per k by sampling with replacement, performing k-means, and computing pairwise Adjusted Rand Index (ARI) values. The median ARI per k was used as the stability score. A noise floor was defined as the 75th percentile of all stability scores; the maximal k above the noise floor (k = 6) was selected (Supplemental Figure 4B). K-means clustering was then run using k = 6 for all larvae. Cluster quality was assessed by comparing temporal profiles of 100 randomly selected pixels to cluster averages. Clusters were color-coded by time-to-peak (fastest in pink, slowest in blue). XY coordinates of spatial maps were stored for cross-larval comparisons.

###### Spatial overlap quantification

Grayscale mask images representing time-to-peak rankings were normalized (Mi_norm = Mi/255). Overlap intensity maps were computed as 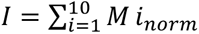. Gaussian smoothing was applied to accommodate minor spatial variability: *I_smooth_*= *gaussian*_*filter*(*I_norm_*,*σ* = 0.8).

###### Kinetics quantification

Cluster fluorescence traces F(t) were converted to ΔF/F using ΔF/F=(F-F_0)/F_0, where F_0 is the baseline fluorescence averaged over a 4 s inactivity period. Time-to-peak T_ifor each cluster i was computed as the time from stimulus onset to the peak of ΔF/F within the recording window. Clusters were ranked by time-to-peak.

###### Statistical analysis

To compare kinetics between clusters, Pearson’s coefficient was used.

###### Sustained activity heatmaps

For sustained activity analyses (400 frames; 80 s post-stimulation), each pixel intensity was normalized by mean activity during the 200 pre-stimulation frames (40 s). For each pixel, the time to decay to half maximum was computed. If half maximum was not reached, the pixel timescale was set to zero. Median pixel timescales across fish were computed per spatial position to generate heatmaps.

###### Overlay with noradrenergic morphology

The fastest cluster (Cluster 1) and the sutained activity cluster overlap map was generated across sessions. A z-stack of noradrenergic neuron morphology was acquired and the plane corresponding to glial imaging was selected. Both datasets were registered to the same internal reference brain and overlaid in Fiji to visualize spatial correspondence.

###### Statistical analysis

Kolmogrov-Smirnoff analysis was used to compute the statistical significance of the different histograms.

#### PERIPHERAL ORGAN MONITORING

Larvae were mounted laterally in low-melting-point agarose (3% for heartbeat quantification; 1.5% for peripheral response visualization), paralyzed via muscle injection of α-bungarotoxin (Tocris), bathed in E3, and placed on a spinning disk microscope equipped with a 5× objective (3i, Denver, USA). Animals were illuminated with continuous IR backlight (890 nm) and received intermittent blue light stimulation (466 nm; 50 ms pulse width; 10 Hz; 3 s total). Stimulation onset was triggered by a TTL pulse generated by a digitizer (Digidata 1440A, Axon Instruments) controlled by Clampex v10.3 (Axon Instruments). Cardiac activity was recorded at 300 fps for 60 s using RD Vision software with an IDS camera (iDS-UI-3060CP-M-GL R2). A 600 nm long-pass filter was placed in the detection path to prevent blue-light absorption.

##### Analysis

For heartbeat quantification, TIFF image sequences were processed in Fiji. A circular ROI was drawn over the pericardium to extract pixel intensity over time. Background fluctuations were accounted for using an ROI placed outside the fish. A custom Python pipeline performed (1) background subtraction, (2) spectral analysis using scipy.signal to compute a spectrogram by Fourier transformation, (3) bandpass filtering to isolate oscillations < 5 Hz, (4) oscillation counting via a zero-crossing method, and (5) sliding-window analysis using a 3s window and 1s step size to compute oscillation count and instantaneous frequency (Hz). For pre–stim–post analysis, mean instantaneous frequency was computed over 3s before stimulation, 3 s during stimulation, and 3 s after stimulation. Instantaneous heart rate (Hz) was estimated via zero-crossing counts in bandpass-filtered signals (<5 Hz) using a 3s sliding window with 1s step. Pre–stim–post values were computed from 3s windows before, during, and after stimulation

##### Statistical analysis

Wilcoxon signed-rank test was used to compare the heartbeat metrics across three points. For peripheral response visualization, standard deviation projections of image sequences acquired 3 s before, during, and after stimulation were computed to visualize motion.

#### ELECTRICAL STIMULATIONS

Electrical stimulation was performed like previously described (Carbo-Tano, Lapoix et al 2023). At 6 dpf, larvae were embedded in 3% low-melting agarose (Thermo Fisher Scientific) in a 35-mm Petri dish filled with external bath solution (134 mM NaCl, 2.9 mM KCl, 2.1 mM CaCl₂·H₂O, 1.2 mM MgCl₂, 10 mM glucose, 10 mM HEPES; pH 7.4; 290 mOsm). Agarose was removed caudal to the swim bladder. Glass-coated tungsten microelectrodes (0.7–3.1 MΩ; 1–2 μm tip) were used for monopolar stimulation in the obex region, positioned at 35° using a motorized micromanipulator (MP-285A, Sutter Instruments). Stimulation with 3-min rest periods was delivered via an isolated stimulation unit triggered by Digidata 1440A controlled by Clampex v10.3. The electrode position was adjusted to identify sites eliciting swimming bouts. Trials lasted 1 min using 2-ms negative pulses (10 Hz, 0.1–3 μA, 40 s) with 2-min rest periods. Trials included electrical stimulation alone or electrical stimulation combined with optogenetic activation (3s, 10Hz, 50ms pulse width of 460nm) LED through the condenser; Digidata 1440A; Clampex v10.3).

#### VENTRICLE INJECTIONS

Hindbrain ventricle injections were performed using a Picospitzer device (World Precision Instruments) and a fine glass needle. Larvae were injected with vehicle aCSF (in mM: 134 NaCl, 2.9 KCl, 1.2 MgCl₂, 10 HEPES, 10 glucose, 2.1 CaCl₂; 290 mOsm·kg⁻¹; pH 7.7–7.8 adjusted with NaOH). Norepinephrine hydrochloride (Sigma, A7256) was diluted to different concentrations in aCSF. Noradrenaline (0.5 nL) was injected at concentrations ranging from 7 nM, 35nM, 140nM and 3 mM. Final ventricular concentrations were estimated and reported based on rhombencephalic ventricular volume measurements (3.5nL estimated, Fame et al., 2016), yielding an estimated range of ∼1 nM, 5nM, 20nM and ∼430 μM respectively. Prazosin was hydrochloride (Tocris; 0623) diluted to 50 μM in aCSF to an approximate final ventricular concentration of ∼7.1 μM. Injection volumes were calibrated to 0.5 nL and the successful ventricular fill was validated by co-injecting 40 μM Alexa 547(Invitrogen, A10438).

#### IMMUNOHISTOCHEMISTRY

6 dpf larvae were euthanized in 0.2% MS-222 (Sigma-Aldrich) in system water. Larvae were fixed in PBS (18912-014, Gibco) containing 4% PFA (15714, Electron Microscopy Sciences), 1% DMSO (D8418, Sigma-Aldrich), and 0.3% Triton X-100 (T9284, Sigma-Aldrich) overnight at 4°C. Samples were rinsed once in 0.3% PBST. Larvae were permeabilized in acetone for 10 min at −20°C and washed three times for 10 min in 0.3% PBSTx. For experiments requiring improved penetration, skin, eyes, jaw, and yolk were removed prior to staining. Samples were blocked in 0.1% Bovine Serum Albumin (Sigma, A4503- 100G) in 0.3% PBSTx at room temperature. Primary antibodies were diluted in blocking solution and incubated overnight at 4°C, including chicken anti-GFP (1:400; ab13970; Abcam), mouse GT335 (1:400; A40251903; AdipoGen life sciences), and anti-adra1a (1:200; UJ290225; Invitrogen). Samples were washed three times for 1 h in 0.3% PBSTx. Alexa Fluor–conjugated secondary antibodies were diluted in blocking solution and incubated overnight at 4°C together with DAPI (0.1%). Secondary antibodies were used at 1:400 and included Alexa Fluor 568 goat anti-rabbit IgG (A11012; Invitrogen), Alexa Fluor 488 goat anti-chicken IgG (A11039; Invitrogen), and Alexa Fluor 647 goat anti-mouse IgG (A32728; Invitrogen). Samples were washed three times for 1 h in 0.3% PBSTx and stored in PBS at 4°C. Zebrafish larvae were mounted dorsally in Mounting Medium (Ibidi, 50001) and imaged on an inverted SP8 DLS confocal microscope (Leica) or a spinning disk confocal microscope (3i, Intelligent Imaging Innovations) with 20× or 40× water-immersion objectives (NA 1.0). Image acquisition, stitching, and processing were performed using Fiji^111^.

## Data and code availability

All python scripts will be made available publicly post publication of the manuscript at (https://github.com/wyartlab/Dhanasekar_et_al_Cell_Press).

## Author contributions

M.D.: Conceptualization, Methodology, Investigation, Formal analysis, Visualization, Writing – original draft, Writing – review & editing.

K.F.: Methodology and Investigation of *in vivo* patch clamp recordings (input resistance, excitability tests, synaptic connectivity investigation) and ventral nerve root recordings in rostral segments. Analysis- inputs for conceptualization of analysis, inputs on Visualization, Writing – review & editing.

E.T.L.: Methodology and Investigation of ventral nerve root recordings in medial segments. Analysis- inputs for conceptualization of analysis, Writing – review & editing.

M.C.T.: Methodology, and Formal investigation, and training M.D. in performing electrical stimulation experiments. Advised on troubleshooting of experimental and analysis protocols. Visualization, Writing – review & editing.

M.G.: Formal analysis – Performed analysis for artefact removal, neuronal recruitment of neurons upon sustained optogenetic activation of noradrenergic neurons and registration of recruitment maps to known transgenic lines. Performed sustained glial activity analysis. Writing – review & editing.

C.D.: Investigation – performed *in vivo* loose patch recordings of noradrenergic neurons in the medulla oblongata.

L.M.: Conceptualization, Analysis, Methodology – k-means clustering algorithm for glial calcium activity.

T.M. & A.W.: Supervision of M.G., Conceptualization – Artefact removal for neuronal recruitment. Writing – review & editing.

C.W.: Conceptualization, Methodology, Formal analysis, Visualization, Writing – original draft, Writing – review & editing, Funding acquisition.

## Conflict of interest

None of the authors declare no conflict of interest.

## Funding

This work was supported by Fondation Bettencourt Schueller FBS-don-0031 to C.W. and FBS-don-1364 to T.M., the European Research Council ERC Consolidator ERC-CoG-Exploratome #101002870, the team grant Fondation pour la Recherche Médicale FRM-EQU202003010612, the Agence Nationale pour la Recherche (ANR) via the ANR-22-CE37-0023 LOCOCONNECT (supporting M.G.), ANR-23-CE16-0017-02 RocSMAP, ANR-21-CE13-0008 ASCENTS, ANR-21-CE14-0042 MOTOMYO, ANR-24-CE16-7992 CIRCOLOCO. C.W was also awarded European Union’s Horizon 2020 Research and Innovation program under the Marie Skłodowska-Curie Grant No. 813457 (https://zenith-etn.com) and the Richard Mille Fund (RMF) grant that she coordinates at the Paris Brain Institute (Institut du Cerveau, https://parisbraininstitute.org/). MD was funded for three years through ED3C doctoral school fellowship. K.F. received support from the Richard Mille Fund (RMF) via the Deep Brain Stimulation project of the Paris Brain Institute. E.T.L. was supported by the Human Frontier Science Program (HFSP-LFP) LT-0022/2023. M.C.T. received support from the Prestige postdoctoral fellowship of the city of Paris.

## Acknowledgments

We thank Antoine Arneau, Camille Lejeune, Nolwenn Jezequel and Sophie Nunes Figueiredo, from Phenoparc for fish care. We thank Dr. Olivier Mirat for his input with tail tracking, Pierre Tissierand the entire FabLab for building the freely swimming optogenetic setup. Dr. Misha Ahrens for sharing published transgenic fish lines. We thank Dr. Ariel Levine and Dr. Giulia Messa for their precious input on the manuscript, Dr. Gautam Sridhar and Dr. Antonio Carlos Costa for their input on the clustering algorithm. We thank Olivier Renaud, Claire Lovo from the ICM.Quant core facility for guidance on optical setups. We thank all the members of the Wyart lab “SIBBIL” (Sensorimotor Integration Brain Body Interaction Lab, https://wyartlab.org) for their input and feedback throughout the study.

---

**Figure S1.**
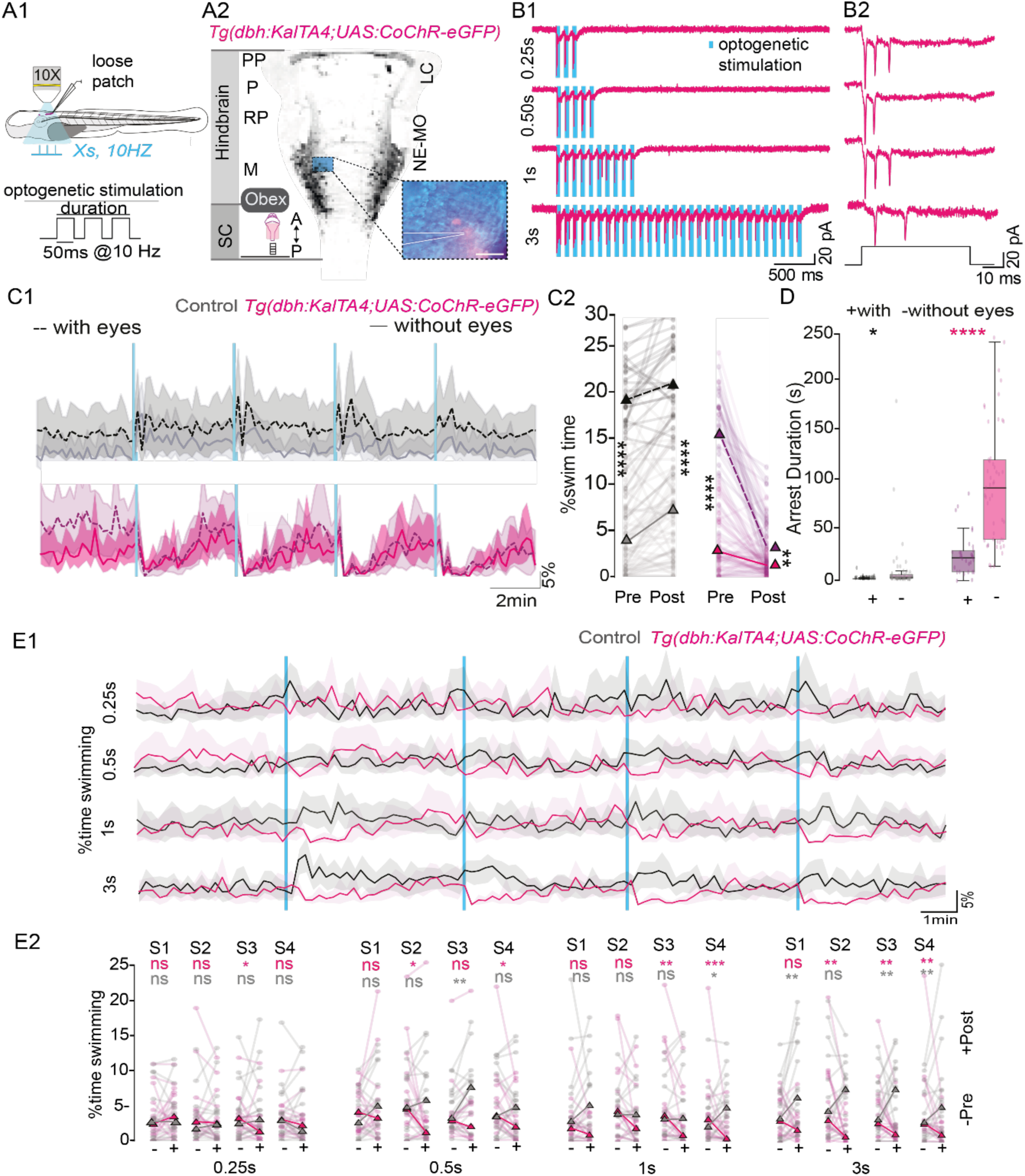
Electrophysiological calibration of optogenetic activation of noradrenergic neurons, effect of enucleation and trial number, Related to Figure 1. **(A1-A2)** Experimental setup for performing targeted loose patch recordings from a noradrenergic neuron located in the medulla oblongata and expressing CoChR-eGFP. SC, spinal cord; PP, prepontine; P, pontine; RP, retropontine; M, medulla; A, anterior, P, posterior. Scale bar, 100 μm. **(B1)** Loose-patch voltage-clamp recordings from medullary noradrenergic neurons in *Tg(dbh:KalTA4;UAS:CoChR-eGFP)* larvae confirm that 50 ms-long light stimulation reliably elicits 1-3 action potentials. Representative traces show spike responses to 10 Hz blue light trains of varying durations (0.25s, 0.5s, 1 s, and 3 s; n = 1 neuron). **(B2)** Zoomed traces to visualize the spikes elicited per light pulse. **(C1)** Locomotor activity computed as a fraction of time spent swimming per time bin of 10s with and without enucleation for 3s-long train stimulation in *Tg(dbh:KalTA4;UAS:CoChR-eGFP)* larvae and control siblings shows comparable decrease in locomotor activity in both enucleated and in animals with eyes. **(C2)** Fraction of time spent swimming computed for 120s-long time bin pre and post onset of stimulus in control and opsin expressing animals with and without eyes. (n= 11 control and 11 transgenic larvae, 4 values per larva for 3s long train stimulation duration). Mann- Whitney U statistical test was performed. Pairwise comparisons of control animals with eyes and without eyes in pre stimulus condition p = 1.94 x 10-19, opsin +ve animals with and without eyes in post stimulus condition p = 8.60 x10-8. Pairwise comparisons of control animals with eyes and without eyes in post stimulus condition p = 2.51 x 10-19, opsin-expressing larvae with and without eyes in post stimulus condition:0.00245. **(D)** Arrest duration comparisons between *Tg(dbh:KalTA4;UAS:CoChR-eGFP)* larvae and control siblings with and without eyes (t-test pairwise comparisons between control animals with and without eyes p = 0.040, and opsin-expressing larvae, p = 3.18×10-9). **(E1)** Locomotor activity represented as a fraction of swimming time in a time bin of 10s for different durations of light pulse shows robust induction of motor arrest in durations for four trials. **(E2)** Fraction of time spent swimming computed for time bin of two minutes before and after stimulus onset computed for each trial (S1-S4) individually for the different stimulation durations. Mann- Whitney U statistical test was performed to evaluate the statistical significance of the data. The p values are presented in Supplementary Table 1 (n = 16 control and 16 transgenic, 4 values per larva for an optical train stimulation duration, 4 stimulations per larva for a given optical train duration).

**Figure S2.**
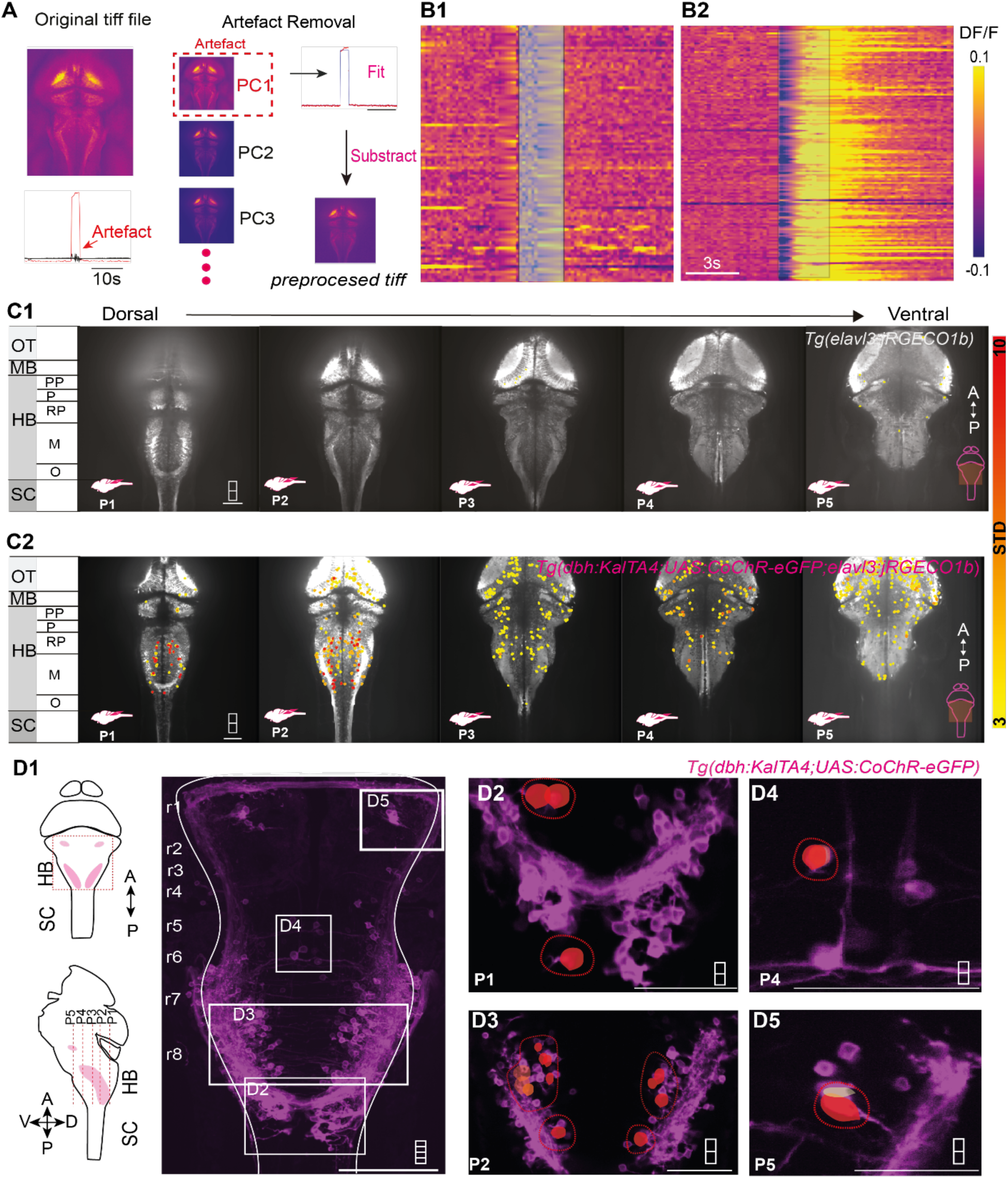
Optogenetic recruitment of noradrenergic neurons in Tg(dbh:KalTA4;UAS:CoChR-eGFP) larvae versus control sibling, Related to Figure 2. **(A)** Detailed pipeline for artifact removal due to blue light absorption of the jRGECO1b sensor in the preprocessing step. **(B1–B2)** Raster plots showing the activity of detected neurons in the control sibling (B1) and *Tg(dbh:KalTA4;UAS:CoChR-eGFP)* larva (B2). **(C1)** Recruitment upon blue light illumination in enucleated control sibling shows minimal recruitment of neurons across z-planes from optic tectum to spinal cord. OT, optic tectum; MB, midbrain; HB, hindbrain; SC, spinal cord; P, pontine; RP, retropontine; M, medulla; A, anterior; P, posterior. Scale bar, 100 μm. **(C2)** Recruitment analysis in enucleated *Tg(dbh:KalTA4;UAS:CoChR-eGFP)* larva shows extensive recruitment of neurons across z-planes from optic tectum to spinal cord. Scale bar, 100 μm. **(D1)** Maximum intensity projection of 6 dpf *Tg(dbh:KalTA4;UAS:CoChR-eGFP)* labelling different subpopulations of noradrenergic neurons. Scale bar,100 μm. **(D2-D5)** Activity-dependent recruitment of noradrenergic neuron subpopulations including area postrema **(D2),** medulla oblongata **(D3),** retropontine cluster **(D4),** and locus coeruleus **(D5)** upon optogenetic stimulation, shown across z-planes. Scale bars,50 μm.

**Figure S3.**
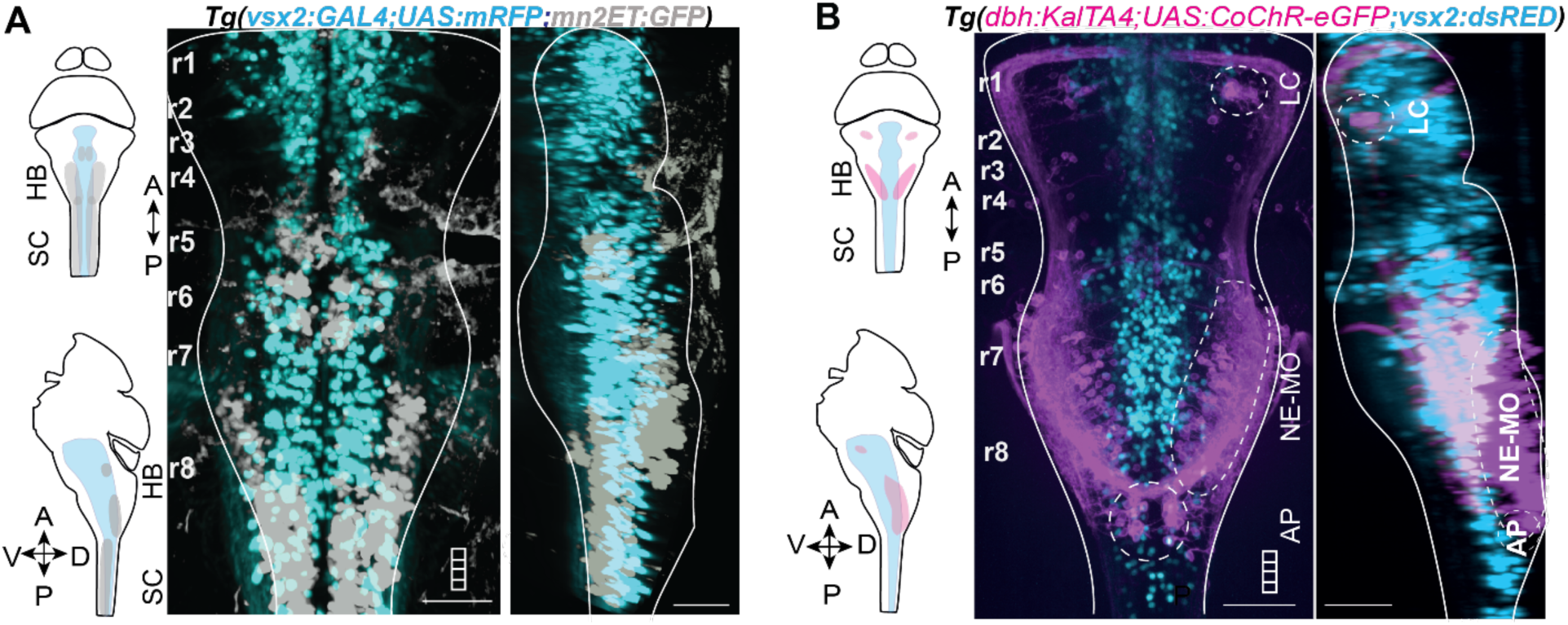
The obex is the site of anatomical convergence of noradrenergic neurons, vagal motor neurons and reticulospinal neurons. Related to Figure 2. **(A)** Noradrenergic neurons are located laterally to the *vsx2+* reticulospinal neurons in the brainstem as shown by the maximum intensity projections of 6 dpf *Tg(dbh:KalTA4;UAS:CoChR-eGFP;vsx2:DsRed)* larval zebrafish brain. Left, dorsal view. Right, lateral view. Noradrenergic neurons, magenta; *vsx2+* reticulospinal neurons, cyan. HB; Hindbrain; SC,spinal cord; P, pontine; RP, retropontine; M, medulla; A, anterior; P, posterior. Scale bar, 50 μm. **(B)** *Mn2ET+* neurons are more dorsolaterally located than the *vsx2+* reticulospinal neurons in the brainstem as shown by the maximum intensity projections of 6 dpf *Tg(vsx2:DsRed;mn2ET:GFP)* larval zebrafish brain. Left, dorsal view. Right, lateral view. *vsx2+* reticulospinal neurons, cyan; motor neurons, gray. Scale bar,50 μm.

**Figure S4.**
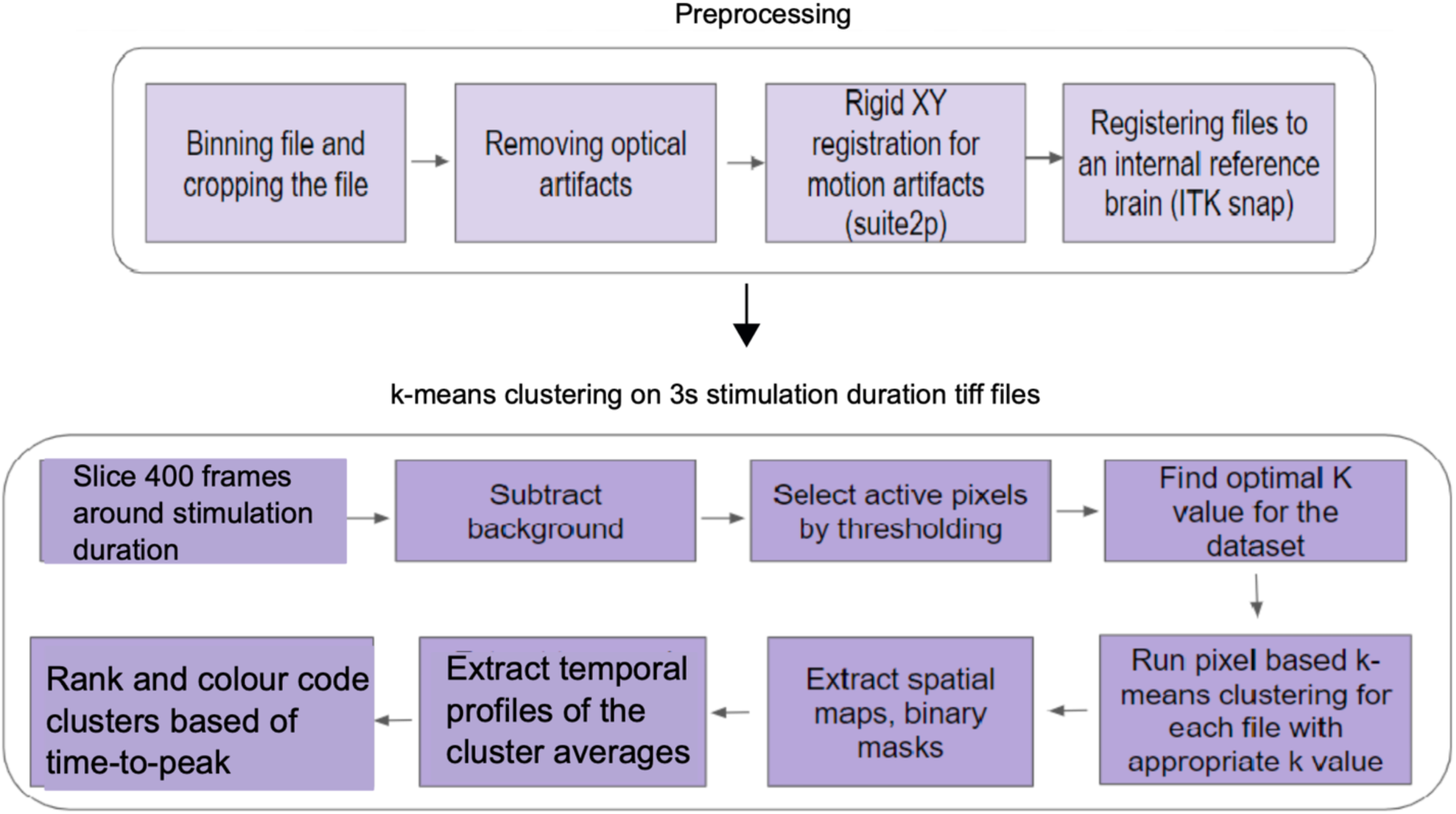
Pipeline used for the pixel-based k-means clustering of glial calcium wave. Related to Figure 5. Flow chart explaining the different steps involved in pixel-based analysis of glial calcium wave. The steps involved three broad categories of preprocessing, k-means clustering on 3s long stimulation duration tiff file and computation of the response rate for all stimulation durations.

**Figure S5.**
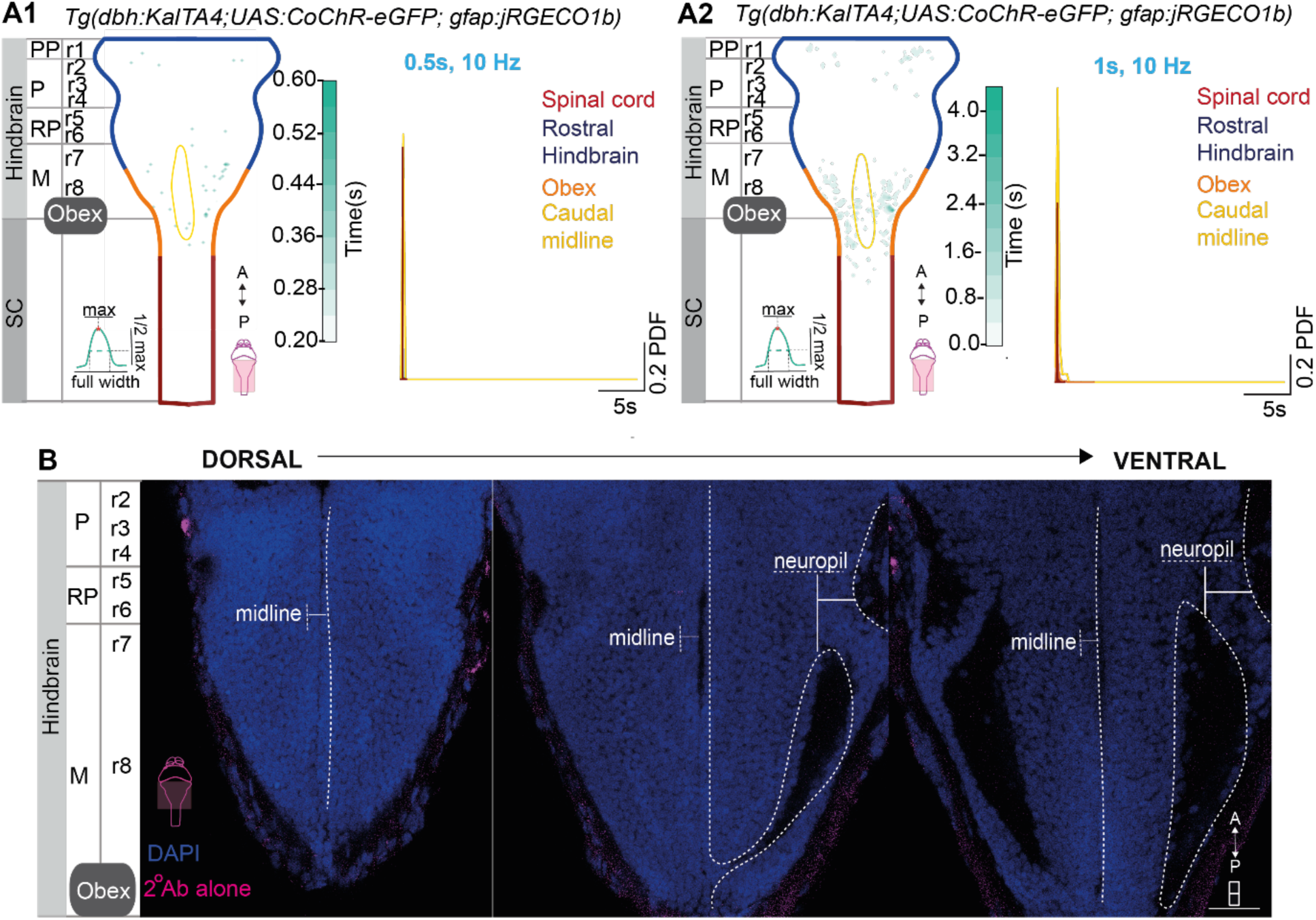
Lack of a widespread glial calcium wave upon short noradrenergic activations. Related to Figure 5 & 6. **(A1)** Spatial map of sustained glial activity for 0.5s reveals lack of consistent hotspots being revealed and lack of activity associated with a singular region. Kolmogorov–Smirnov tests revealed no significant regional differences at 500 ms. Comparisons between rHB and Obex (test statistic = 0.0106, p = 1.0), rHB and SC (test statistic = 0.0014, p = 1.0), and rHB and Midline (test statistic = 0.0047, p = 1.0) were not significant. Similarly, no significant differences were observed between Obex and SC (test statistic = 0.0121, p = 1.0), Obex and Midline (test statistic = 0.0060, p = 1.0), or Midline and SC (test statistic = 0.0061, p = 1.0). rHB, rostral hindbrain; SC, spinal cord; PP, prepontine; P, pontine; RP, retropontine; M, medulla; A, anterior; P, posterior. **(A2)** Spatial map of sustained glial activity for 1s reveals an emerging hotspot at the obex (left) with a significant difference in the duration of sustained activity in the obex region (right). Kolmogorov–Smirnov tests revealed significant regional differences at 1 s. Comparisons showed significant differences between rHB and Obex (test statistic = 0.0781, p = 7.7 × 10⁻⁴), rHB and SC (test statistic = 0.0581, p = 9.9 × 10⁻⁴), and rHB and Midline (test statistic = 0.1697, p = 2.7 × 10⁻³). Significant differences were also observed between Obex and SC (test statistic = 0.1361, p = 5.2 × 10⁻¹³) and between Midline and SC (test statistic = 0.2278, p = 9.2 × 10⁻⁶). No significant difference was detected between Obex and Midline (test statistic = 0.0917, p = 0.31). **(B)** Single plane images of different planes throughout the brain show no intensity for secondary antibodies alone revealing the specificity of the enrichment. Scale bar, 50 μm. A, anterior; P, posterior; P, pontine; RP, retropontine; M, medulla.

**Figure S6.**
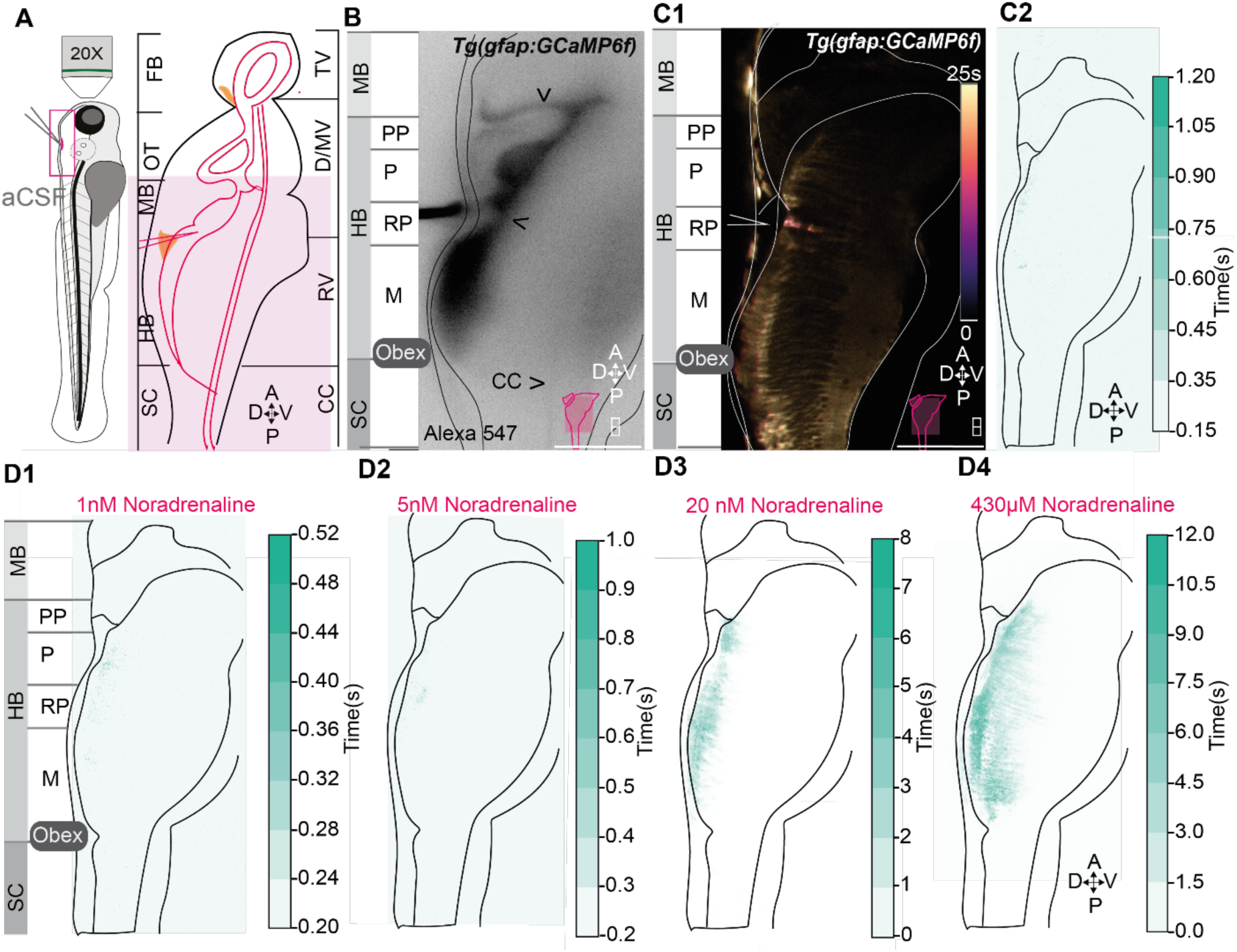
Calibration of ventricular injection, control injection of aCSF, Related to Figure 7. **(A)** Experimental paradigm showing the site of ventricular injection of aCSF in the rhombencephalic ventricle. FB, forebrain; OT, optic tectum; MB, midbrain; HB, hindbrain; SC, spinal cord; TV, telencephalic ventricle; D/MV, diencephalic/mesencephalic ventricle; RV, rhombencephalic ventricle; CC, central canal; A, anterior; P, posterior; D, dorsal; V, ventral. **(B)** Visualization of successful ventricular injection through 40μM Alexa 547 dye shows the filling of the ventricular system with the dye. **(C1)** Representative timeseries color-coded based on time-to-peak upon injection of aCSF in the rhombencephalic ventricle reveals a lack of glial calcium wave induction in *Tg(gfap:GCaMP6f)* larvae. (C2) Spatio-temporal map of the sustained glial activity reveals minimal activity. **(D1–D4)** Noradrenaline induces concentration-dependent calcium responses in obex ependymal glia. Representative images following application of 1 nM (D1), 5 nM **(D2),** 20 nM **(D3),** and 430 μM **(D4)** noradrenaline. Low concentrations (1 nM, 5 nM) elicit minimal calcium activity. Higher concentrations (20 nM, 430 μM) trigger sustained glial calcium responses, with ventricular-contacting processes emerging as a long-lasting component.

## Supplementary Videos

**Supplementary Video 1**.

Loss of dorsoventral orientation of the *Tg(dbh:KalTA4;UAS:CoChR-eGFP)* upon 3s optogenetic stimulation of noradrenergic neurons. Recorded at 100Hz, playback rate 1Hz.

**Supplementary Video 2.**

Whole-body responses observed in *Tg(dbh:KalTA4;UAS:CoChR-eGFP)* upon 3s optogenetic stimulation of noradrenergic neurons. Recorded at 100Hz, playback rate of 7Hz.

**Supplementary Video 3.**

Z stack of the 5 dpf *Tg(dbh:GAL4; UAS:CoChR-YFP)* transgenic larva showing from a dorsal view on the brainstem noradrenergic projections from medulla oblongata and area postrema respectively reaching the obex and ventricular midline. Rostral, up.

**Supplementary Video 4.**

Time series showing the glial calcium wave upon injection of 430μM noradrenaline in the rhombencephalic ventricle. Acquisition at 5Hz, playback at 10Hz.

## Notes

### Competing Interest Statement

The authors have declared no competing interest.

