## Supplemental Table 1 for "Noradrenaline prolongs the state of motor arrest via cerebrospinal fluid signaling"

### Supplementary table 1: Statistical comparison of the trial wise computation of fraction of time spent swimming in control siblings and transgenic larvae related to Figure S1E2.

| Comparison percentage swimming | Stim | Statistic | p-value | Significance | Duration |
|---|---|---|---|---|---|
| Pre Control vs Experimental | 1 | 124 | 0.894242 | ns | 0.25s |
| Post Control vs Experimental | 1 | 123 | 0.865202 | ns | 0.25s |
| Control Pre vs Post | 1 | 32 | 0.345448 | ns | 0.25s |
| Experimental Pre vs Post | 1 | 55 | 0.528168 | ns | 0.25s |
| Pre Control vs Experimental | 2 | 104 | 0.375654 | ns | 0.25s |
| Post Control vs Experimental | 2 | 127.5 | 1 | ns | 0.25s |
| Control Pre vs Post | 2 | 48 | 0.495521 | ns | 0.25s |
| Experimental Pre vs Post | 2 | 45 | 0.252228 | ns | 0.25s |
| Pre Control vs Experimental | 3 | 113.5 | 0.597712 | ns | 0.25s |
| Post Control vs Experimental | 3 | 178 | 0.062001 | ns | 0.25s |
| Control Pre vs Post | 3 | 27 | 0.060893 | ns | 0.25s |
| Experimental Pre vs Post | 3 | 25 | 0.024963 | * | 0.25s |
| Pre Control vs Experimental | 4 | 128 | 1 | ns | 0.25s |
| Post Control vs Experimental | 4 | 133 | 0.865079 | ns | 0.25s |
| Control Pre vs Post | 4 | 57 | 0.864705 | ns | 0.25s |
| Experimental Pre vs Post | 4 | 42 | 0.19281 | ns | 0.25s |
| Pre Control vs Experimental | 1 | 100 | 0.299817 | ns | 0.5s |
| Post Control vs Experimental | 1 | 138 | 0.72031 | ns | 0.5s |
| Control Pre vs Post | 1 | 22 | 0.015503 | * | 0.5s |
| Experimental Pre vs Post | 1 | 59 | 0.668549 | ns | 0.5s |
| Pre Control vs Experimental | 2 | 122 | 0.835636 | ns | 0.5s |
| Post Control vs Experimental | 2 | 179 | 0.057003 | ns | 0.5s |
| Control Pre vs Post | 2 | 45 | 0.252228 | ns | 0.5s |
| Experimental Pre vs Post | 2 | 21 | 0.013092 | * | 0.5s |
| Pre Control vs Experimental | 3 | 139.5 | 0.678423 | ns | 0.5s |
| Post Control vs Experimental | 3 | 194 | 0.013529 | * | 0.5s |
| Control Pre vs Post | 3 | 14 | 0.003357 | ** | 0.5s |
| Experimental Pre vs Post | 3 | 57 | 0.596588 | ns | 0.5s |
| Pre Control vs Experimental | 4 | 125 | 0.924933 | ns | 0.5s |
| Post Control vs Experimental | 4 | 173.5 | 0.089591 | ns | 0.5s |
| Control Pre vs Post | 4 | 68 | 1 | ns | 0.5s |
| Experimental Pre vs Post | 4 | 27 | 0.033539 | * | 0.5s |
| Pre Control vs Experimental | 1 | 141.5 | 0.404821 | ns | 1s |
| Post Control vs Experimental | 1 | 187.5 | 0.008023 | ** | 1s |
| Control Pre vs Post | 1 | 23 | 0.064039 | ns | 1s |
| Experimental Pre vs Post | 1 | 34 | 0.245494 | ns | 1s |
| Pre Control vs Experimental | 2 | 112 | 0.766785 | ns | 1s |
| Post Control vs Experimental | 2 | 150 | 0.242578 | ns | 1s |
| Control Pre vs Post | 2 | 47 | 0.729891 | ns | 1s |
| Experimental Pre vs Post | 2 | 32 | 0.065399 | ns | 1s |
| Pre Control vs Experimental | 3 | 109.5 | 0.692603 | ns | 1s |
| Post Control vs Experimental | 3 | 162.5 | 0.095697 | ns | 1s |
| Control Pre vs Post | 3 | 38 | 0.229309 | ns | 1s |
| Pre Experimental Pre vs Post | 3 | 9 | 0.006319 | ** | 1s |
| Post Control vs Experimental | 4 | 106 | 0.593595 | ns | 1s |
| Control vs Experimental | 4 | 199 | 0.001908 | ** | 1s |
| Control Pre vs Post | 4 | 23 | 0.035339 | * | 1s |
| Experimental Pre vs Post | 4 | 7 | 0.00058 | *** | 1s |
| Pre Control vs Experimental | 1 | 128 | 1 | ns | 3s |
| Post Control vs Experimental | 1 | 205 | 0.003937 | ** | 3s |
| Control Pre vs Post | 1 | 14 | 0.003357 | ** | 3s |
| Experimental Pre vs Post | 1 | 35 | 0.093445 | ns | 3s |
| Pre Control vs Experimental | 2 | 147 | 0.48561 | ns | 3s |
| Post Control vs Experimental | 2 | 222 | 0.000425 | *** | 3s |
| Control Pre vs Post | 2 | 55 | 0.528168 | ns | 3s |
| Experimental Pre vs Post | 2 | 22 | 0.015503 | * | 3s |
| Pre Control vs Experimental | 3 | 130 | 0.954913 | ns | 3s |
| Post Control vs Experimental | 3 | 213 | 0.001434 | ** | 3s |
| Control Pre vs Post | 3 | 31 | 0.057678 | ns | 3s |
| Experimental Pre vs Post | 3 | 6 | 0.002162 | ** | 3s |
| Pre Control vs Experimental | 4 | 124 | 0.895007 | ns | 3s |
| Post Control vs Experimental | 4 | 200.5 | 0.006606 | ** | 3s |
| Control Pre vs Post | 4 | 20 | 0.023096 | * | 3s |
| Experimental Pre vs Post | 4 | 12 | 0.006406 | ** | 3s |
