## Supplemental Table 2 for "Noradrenaline prolongs the state of motor arrest via cerebrospinal fluid signaling"

**Supplementary Table 2: Wilcoxon test comparisons of the action potential parameters related to Figure 3, J-N. Parameter**

| Parameter | **Comparison** | **Statistic** | **p-value** | **Significance** |
|---|---|---|---|---|
| Amplitude | Pre Light vs. Pre No Light | 0 | 0.25 | ns |
| Amplitude | Post Light vs. Post No Light | 3 | 1 | ns |
| Amplitude | Pre vs. Post Light | 3 | 1 | ns |
| Amplitude | Pre vs. Post No Light | 0 | 0.25 | ns |
| Frequency | Pre Light vs. Pre No Light | 2 | 0.75 | ns |
| Frequency | Post Light vs. Post No Light | 2 | 0.75 | ns |
| Frequency | Pre vs. Post Light | 2 | 0.75 | ns |
| Frequency | Pre vs. Post No Light | 3 | 1 | ns |
| ISI | Pre Light vs. Pre No Light | 1 | 0.5 | ns |
| ISI | Post Light vs. Post No Light | 0 | 0.25 | ns |
| ISI | Pre vs. Post Light | 2 | 0.75 | ns |
| ISI | Pre vs. Post No Light | 2 | 0.75 | ns |
| Latency | Pre Light vs. Pre No Light | 0 | 0.25 | ns |
| Latency | Post Light vs. Post No Light | 1 | 0.5 | ns |
| Latency | Pre vs. Post Light | 3 | 1 | ns |
| Latency | Pre vs. Post No Light | 0 | 0.25 | ns |
